# Systematic comparisons between long-read and short-read based amplicon sequencing to characterize mixed microalgal communities

**DOI:** 10.64898/2026.04.17.719029

**Authors:** Zihan Dai, Md Mahbubul Alam, Benjamin Gincley, Farhan Khan, Ga-Yeong Kim, Hannah Molitor, Jeremy Guest, Ian M. Bradley, Ameet J. Pinto

## Abstract

The 18S rRNA gene has emerged as the primary molecular marker for amplicon-based characterization of microalgal communities, including in wastewater treatment systems, yet trade-offs between short- and long-read approaches remain poorly defined. We systematically compared V8–V9 short-read sequencing (Illumina MiSeq), full-length long-read sequencing with ss5ss3 primers (PacBio Sequel II), and computationally extracted V8–V9 regions from long-read data. Both *in silico* and *in vitro* analyses confirmed V8–V9 captured broader taxonomic coverage than ss5ss3, though partial reference sequences and taxonomic mis-annotations biased *in silico* assessments. Long-read’s taxonomic advantage was database-dependent, constrained by SILVA databases genus-level curation but fully realized when paired with the species-level-curated and eukaryotes-focused PR² (Protist Ribosomal Reference) database. Long-read sequencing uniquely identified amplicon sequence variants (ASVs) assigned to key phosphorus assimilators (*Scenedesmus obliquus*, *Desmodesmus* sp., and *Acutodesmus* sp.) at species level during successful phosphorus removal in a full-scale microalgal cultivation system, while V8–V9 short-read sequencing revealed ASVs assigned to algal-predatory (Leptophryidae) and bacterivorous (Choanoflagellata and Rhogostoma-lineage) protists when performance declined, suggesting grazing pressure on the phosphorus-removing community. Although both approaches performed comparably for operational monitoring, these complementary strengths support short-read sequencing for routine community profiling and long-read sequencing for detailed functional investigations of Chlorophyta.

## Introduction

Microalgal-based wastewater treatment systems provide effective nutrient removal from wastewater while potentially generating valuable biomass for biofuels, biopolymers, and fertilizers.^1–5^ These systems can remove nitrogen and phosphorus below the current limit-of-technology levels via nutrient assimilation which complements the conventional nutrient removal processes of nitrification/denitrification and luxury phosphorus uptake by bacteria.^6,7^ Utilizing microalgal monocultures for wastewater treatment is impractical due to the inherent presence of diverse microorganisms. Instead, microalgal processes for wastewater treatment primarily rely on mixed communities of diverse algal species.^8^ Understanding the composition and structure of these mixed microalgal communities is critical for optimizing treatment efficiency, as different species may exhibit varying nutrient removal capabilities.^8–11^ Insights into community composition can also advance strategies for robust process control by elucidating links between microalgal community dynamics and process performance and stability.^12–16^ Further, species composition directly affects biomass quality and harvesting efficiency, which has a direct impact on resource recovery potential.^17,18^

Traditional methods for studying microalgal communities rely on microscopy for species identification and enumeration.^19,20^ Although microscopy provides valuable morphological details, its application is constrained by the need for extensive taxonomic expertise, significant time investment, and the inability to identify organisms without distinct morphological cues. High-throughput sequencing methods have revolutionized microalgal community analysis through morphology-independent species identification and rapid parallel sample processing. These advances have made comprehensive community analysis more efficient and accessible.^21,22^ Researchers have employed various genetic markers for microalgal community characterization, including the 18S rRNA gene,^22–24^ internal transcribed spacers (ITS1 and ITS2),^25,26^ mitochondrial cytochrome c oxidase subunit I ^27,28^, the *tufA* gene,^29^ and the *rbcL* gene.^30^ The 18S rRNA gene, specifically the V4 and V8–V9 hypervariable regions, has emerged as the primary target for microalgal community analysis due to its broad taxonomic coverage and due to the presence of well-curated reference SSU rRNA gene databases.^21^ Using improved primers targeting the V4 and V8–V9 regions, Bradley *et al.* found that shorter solids retention times promote more diverse communities dominated by filamentous eukaryotes, while longer retention times support stable communities of unicellular and colonial green algae.^31^ Jo *et al.* used V8–V9 hypervariable region sequencing to note the consistently high abundance of *Desmodesmus* sp. (62–74%) in open pond raceways despite changes in multiple environmental variables.^32^ Similarly targeting the V8–V9 region, Alam *et al.* monitored a full-scale system for nine months to demonstrate that stable performance correlates with low eukaryotic diversity dominated by *Scenedesmus* sp. (55–80%).^12^ Ma *et al.* sequenced the V4 region to show that lower substrate loading rate resulted in increased community diversity in a hybrid algal-bacterial granular sludge system.^33^ In a membrane-coupled high-rate algal pond, Aparicio *et al.* used V4 region sequencing of a system with a high abundance of *Desmodesmus* (30–69%) and *Coelastrella* (11–44%) to show that microalgae biodiversity was not affected by hydraulic retention time (HRT) or influent wastewater quality.^34^

Although short-read sequencing has advanced our understanding of microalgal community structure and dynamics, it faces significant limitations due to read length constraints (up to 550 bp).^35^ These constraints reduce taxonomic resolution and restrict the selection of target-specific primer binding sites.^35–39^ The challenges are amplified in complex environmental communities where closely related organisms exhibit high sequence similarity in the targeted regions. Accurate taxonomic classification is further complicated by incomplete reference databases for understudied unicellular eukaryotes, PCR amplification biases, sequencing errors that generate artificial variants, and the presence of length variations and introns in the 18S rRNA genes of some species.^35^ Long-read sequencing technologies can address these technical gaps by providing comprehensive sequence information in a single read, enabling improved taxonomic resolution for eukaryotic communities.^40^ The sequencing accuracy of long-read platforms has increased, as demonstrated by PacBio’s HiFi reads now matching short-read methods in precision to Illumina.^35,38^ However, the complex library preparation process requires high-quality DNA and multiple purification steps, increasing both processing time and costs resulting in higher per-sample costs as compared to short-read platforms.^38^ As a result, long-read sequencing remains less accessible than short-read methods for large-scale environmental surveys and time-series studies requiring numerous sample analyses.

We conducted a systematic comparison of short- and long-read 18S rRNA gene sequencing to characterize microalgal communities in a full-scale microalgal wastewater treatment system.^6^ Our analysis examines three datasets: (1) short-read sequencing of the V8–V9 region sequenced on the Illumina MiSeq platform, (2) full-length long-read data generated on the PacBio Sequel II platform, and (3) computationally-extracted V8–V9 sequences from long-read PacBio data. Through this comparative study, we assess how different sequencing methods affect insights from microalgal community analysis and determine whether the value proposition of improved taxonomic resolution at the higher cost for long-read sequencing relative to that of short-read sequencing results in actionable, process-relevant insights.

## Materials and Methods

### Primer and Database Selection

Four primer sets targeting the 18S rRNA gene were selected from established literature for *in silico* comparison (**Table 1**). Two primer sets targeting the V4 and V8–V9 regions were chosen for short-read sequencing platforms and these result in ∼420 bp and ∼375 bp-long amplicons, respectively. Two universal eukaryotic primer sets (ss5ss3 and EukAB) were selected for long-read sequencing platforms, targeting nearly the full-length 18S rRNA gene with amplicons of ∼1800 bp.

**Table 1:**
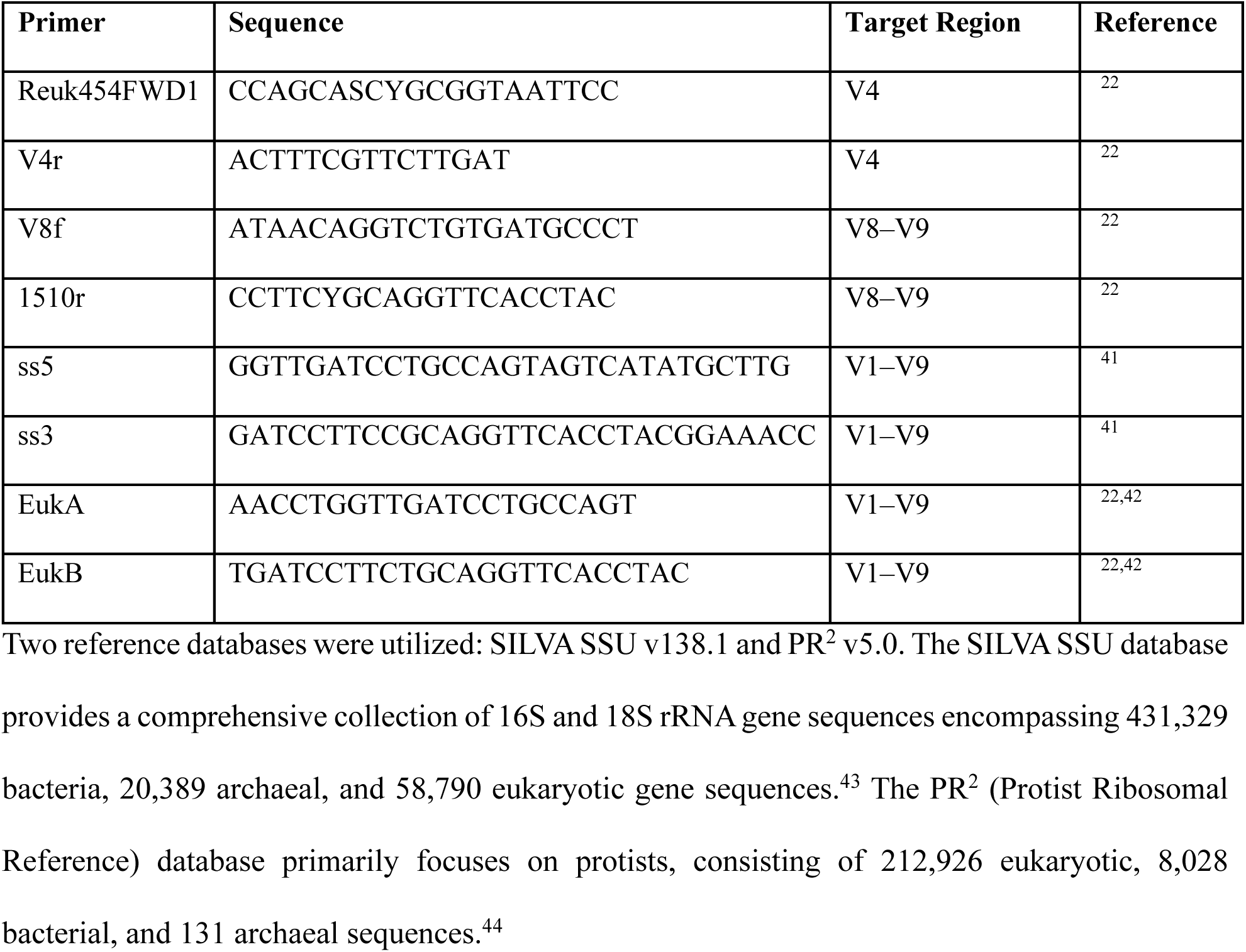
Primers initially selected for the *in silico* comparison between short- and long-read based amplicon sequencing. Target regions range from specific hypervariable regions (V4, V8–V9) to near-complete 18S rRNA gene coverage (V1–V9).

### In silico PCR analysis

*In silico* polymerase chain reaction (PCR) analysis was conducted on rRNA gene sequences from both databases using the following workflow conducted by RESCRIPt within the QIIME2 v2023.5: (1) RNA sequences were reverse transcribed to DNA, (2) sequence with >5 ambiguous base pairs or homopolymer regions ≥8 base pairs were removed and sequences below minimum length thresholds (Archaea: 900 bp, Bacteria: 1200 bp, Eukaryotes: 1400 bp) were discarded, and (3) *in silico* amplicon extraction was conducted using ipcress from exonerate v2.4.0 with four primer pairs listed in Table 1.^45–47^ Following the extraction of *in silico* amplicon sequences, only products generated by both primers (description field in “*forward*” and “*revcomp*”) from ipcress results were retained for further analysis. After evaluating the length distribution for each primer set, EukAB was removed from further consideration due to a wide amplicon length distribution. Outliers for the remaining three primer sets were identified using the Hampel filter (equation 1), where *I* is the interval, and MAD is the median absolute deviation:

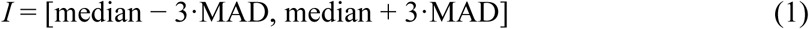

Amplicon sequences outside the following length intervals were removed from the downstream analysis: V4 (410–428 bp), V8–V9 (363–381 bp), and ss5ss3 (1770–1818 bp).

Single primer matching sequences were identified using SeqKit v2.4.0 with the *locate* command.^48,49^ For coverage assessment, hits were deduplicated using reference sequence ID. The full amplicon potential analysis evaluated whether reference sequences could accommodate complete amplification with each primer set. For each primer match, theoretical amplification endpoints were calculated by adding (forward primers) or subtracting (reverse primers) the average amplicon length from the primer binding position. All binding positions were included when multiple matches occurred on a single reference sequence. Reference sequences demonstrated full amplicon potential when calculated endpoints fell within sequence boundaries; specifically, when reference length exceeded the calculated endpoint for forward matches, or when the calculated endpoint remained positive for reverse matches.

### DNA extraction, PCR, and amplicon sequencing

A total of 48 microalgal biomass samples were collected from a full-scale mixed-microalgal wastewater treatment system (EcoRecover; Clearas Water Recovery Inc., U.S.)^6^ between August 10, 2022, and July 18, 2023, as described in detail by Alam *et al.*^12^ DNA extracts were shipped overnight on ice to Georgia Institute of Technology and stored at −20°C until further analysis. Short-read sequencing of the V8–V9 hypervariable region of the 18S rRNA gene was performed in technical duplicate (two independent libraries per DNA extract) on the Illumina MiSeq platform with v3 chemistry (300-cycle paired-end reads) at the Georgia Institute of Technology’s Molecular Evolution Core. For long-read sequencing, the full-length 18S rRNA gene was amplified using ss5ss3 primer pairs in 25 μL reactions using KAPA HiFi HotStart Ready Mix. Cycling conditions consisted of an initial denaturation at 98°C for 30 s, followed by 22 cycles of 98°C for 10 s, 68°C for 30 s, and 72°C for 45 s, with a final extension at 72°C for 120 s. Amplicon library preparation and sequencing were performed by Maryland Genomics following the PacBio barcoded M13 primers and SMRTbell Prep Kit 3.0 protocol on the PacBio Sequel II platform.

### Data processing and statistical analyses

Distinct yet parallel bioinformatic pipelines were employed for the processing of short-read (SR) and long-read (LR) sequencing data. For SR sequences, optimal trimming parameters were initially determined using Figaro to ensure maximal retention of high-quality data.^50^ Subsequently, DADA2 v1.26.0 was utilized for quality trimming, denoising, and merging of sequences into amplicon sequence variants (ASVs).^51^ Further refinement of the dataset was achieved through length filtering, retaining sequences between 305 and 337 base pairs, followed by chimera removal.^51^ Long-read sequences underwent a similar, albeit adapted process. Raw LR sequencing reads were initially filtered based on quality scores, with retention of reads exhibiting a Phred score exceeding Q20. This step was followed by the removal of adapter sequences and primers using Cutadapt v4.4.^52^ ASV inference was again performed using DADA2 v1.26.0, with parameters optimized for LR data: the error estimation function was set to *PacBioErrfun*, the alignment band size (*BAND_SIZE*) was set to 32, and the number of bases for error rate estimation (*nbases*) was set to 2 × 10⁸. Following ASV inference, length filtering was applied, and chimeras were removed with *minFoldParentOverAbundance* set to 8. To generate *in silico* long-to-short (LS) reads, GGTTTCCGTAGGTGAACCTGCGGAAGG was appended to the 3’ end of long-read sequences to account for the length difference between ss5ss3 and V8–V9 reverse primers as appropriate. The V8–V9 amplicons were extracted computationally using ipcress (exonerate v2.4.0),^45^ followed by primer removal with Cutadapt v4.4.^52^

Taxonomic classification was implemented using both SILVA SSU v138.1 and PR^2^ databases. Custom classifiers with SILVA SSU v138.1 database were constructed using the RESCRIPt plugin and quality filtered as outlined in the *in silico* PCR analysis section. The development of the full-length 18S rRNA gene classifier involved: (1) preparing SILVA taxonomy data with species label inclusion, (2) deduplicating filtered sequences and taxonomy data using *’uniq’* mode, and (3) generating the classifier. Additionally, the Ribosomal Database Project (RDP) classifier (*minBoot* = 80) was implemented through DADA2 using the PR^2^ database, which provides more accurate species-level information compared to SILVA. These classification methods were applied across all three datasets (SR, LR, and LS) to enable subsequent comparative analysis of their taxonomic resolution performance.

To standardize the statistical analysis between datasets with different sample structures (i.e., 30 duplicate sequencing libraries per sample for SR data versus 30 single sequencing libraries per sample for LR data), the SR duplicate libraries were combined by summing the ASV counts for each sample while retaining only ASVs shared between duplicate libraries. The datasets were rarefied to standardized sequencing depths: 94,846 reads for SR, 10,893 reads for LR, and 11,306 reads for LS. Data analysis was conducted using R v4.2.1 within RStudio.^53,54^ The data.table and tidyverse packages facilitated data manipulation, while the vegan and phyloseq packages were used for prevalence filtering and rarefaction.^55–58^ Alpha-diversity metrics (observed ASVs, Shannon and Simpson indices) and Bray-Curtis dissimilarity were calculated using phyloseq, with UniFrac distances computed via the phyloseq.extended package.^57,59^ The vegan package was used for Principal Coordinates Analysis (PCoA), distance-based Redundancy Analysis (dbRDA), and Mantel tests.^58^ Pairwise ASV sequence identity was determined using MSA2dist and ape packages.^60,61^ Differential abundance analysis was conducted on samples collected during two periods (August 10, 2022 – November 2, 2022 and December 8, 2022 – April 28, 2023) using ALDEx2.^62–64^ The analysis was performed on raw count data after prevalence and sample size filtering. An effect size threshold of ≥1 and a relative abundance threshold of >1% were applied to identify differentially abundant ASVs. Visualization was implemented through ggplot2, ggpubr, and gghalves packages.^65–67^ The raw sequencing data are available in the NCBI Sequence Read Archive under BioProject accession PRJNA1225776.

## Results and Discussion

### Primer choice is influenced by not only on coverage, but also on resultant PCR product length

*In silico* PCR was performed to assess the theoretical capture efficiency of the four primer pairs against the SILVA and PR^2^ reference databases. The V4 primer pair exhibited the highest coverage, generating 40,281 and 93,308 amplicons from the SILVA and PR^2^ databases, respectively. The V8–V9 primer pair demonstrated moderate efficacy, producing 10,074 and 25,315 amplicons, while the ss5ss3 pair yielded 5,564 and 13,484 amplicons from the respective databases. The EukAB primer pair showed the lowest capture rates, generating 2,360 and 4,833 amplicons. In addition, evaluation of the length distribution of *in silico* PCR-generated amplicons revealed distinct patterns among the primer pairs (Figure 1). V4 and V8–V9 yielded short, tightly distributed amplicons (V4: 431.57 ± 50.43 bp, interquartile range (IQR) = 8 bp; V8–V9: 374.75 ± 42.13 bp, IQR = 5 bp). Despite targeting a substantially longer region, ss5ss3 demonstrated the highest proportional consistency of all primer sets (1797.83 ± 83.30 bp, IQR = 15 bp, coefficient of variation (CV) = 4.63%), suggesting that the targeted region is highly conserved in length across eukaryotic reference sequences. In contrast, EukAB produced the broadest length distribution by every measure (1813.87 ± 208.42 bp, IQR = 47 bp, CV = 11.49%), with significantly greater variance compared to all other primer sets (pairwise Levene’s test, all adjusted p-value < 0.001). Combined with its comparatively low capture efficiency, the excessive length variability of EukAB was considered incompatible with robust downstream analysis, and it was therefore excluded from all subsequent analyses.

**Figure 1:**
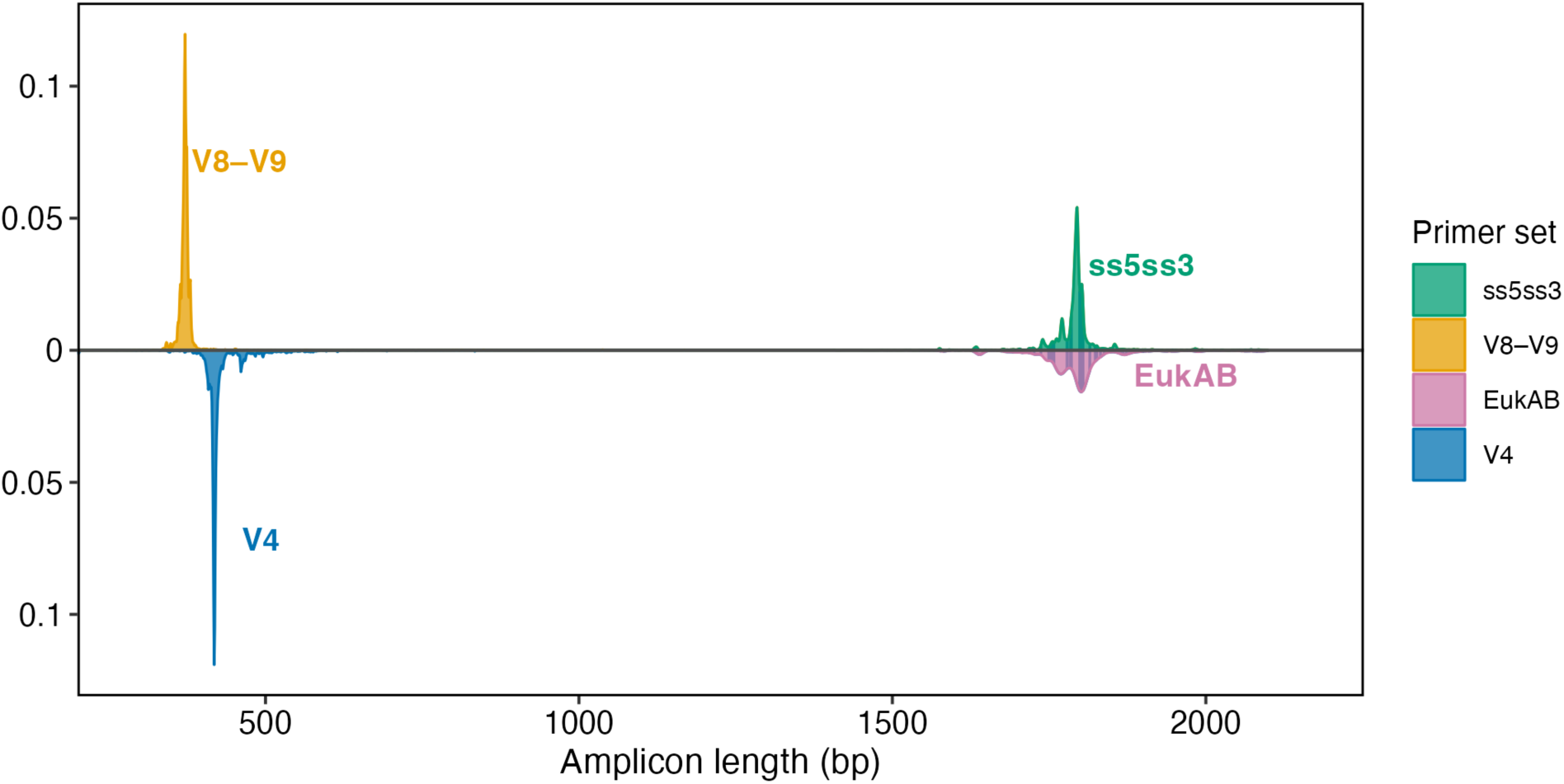
Length distribution of amplicons captured by different primer pairs prior to length filtering. Density curves represent: V4 (blue) and EukAB (reddish) primer pairs (below baseline); V8–V9 (orange) and ss5ss3 (green) primer pairs (above baseline).

To refine the assessment of primer performance, a comprehensive coverage analysis was conducted with the incorporation of a length filtering step to exclude amplicons outside the expected size range for each target region. Amplicons were filtered based on the following ranges: V4 (410–428 bp), V8–V9 (363–381 bp), and ss5ss3 (1770–1818 bp). The coverage analysis of length-filtered amplicons revealed that the V4 primer pair achieved the highest coverage, generating 26,831 (forward = 47,189, reverse = 42,209) and 63,541 (forward = 107,681, reverse = 95,107) amplicons in SILVA and PR^2^ databases, respectively. The V8–V9 primer pair produced 8,610 (forward = 45,602, reverse = 10,524) and 22,109 (forward = 104,546, reverse = 26,226) amplicons, while the ss5ss3 primer pair yielded 4,090 (forward = 13,004, reverse = 8,407) and 10,383 (forward = 24,749, reverse = 21,357) amplicons in SILVA and PR^2^ databases, respectively (Figure 2A). Reverse primers consistently performed poorly as compared to forward primers, with V8–V9_R being substantially limited in reference sequences. The lower capture rates for V8–V9 and ss5ss3 are likely attributable to partial 18S rRNA gene sequences in the databases considering that the V8–V9 targets the gene’s terminal region and ss5ss3 spans nearly the entire gene. To test this hypothesis, full amplicon potential was assessed by adding (for forward strand) or subtracting (for reverse strand) the average amplicon size from the primer binding site position. This calculated position was compared to the reference sequence length to determine if it could accommodate the full amplicon (Figure 2B). V4 primers showed high full amplicon potential (99.89% - 99.99%). For V8–V9, while high potential was shown by the reverse primer (SILVA: 99.89%; PR^2^: 99.68%), much lower potential was exhibited by the forward primer (SILVA: 30.45%; PR^2^: 28.39%). This suggests that in silico amplification failure with V8–V9_R is often due to short reference sequences. Low full amplicon potential (44.33% - 55.67%) was observed for both ss5 and ss3 primers, likely because they target nearly the entire 18S rRNA gene.

**Figure 2:**
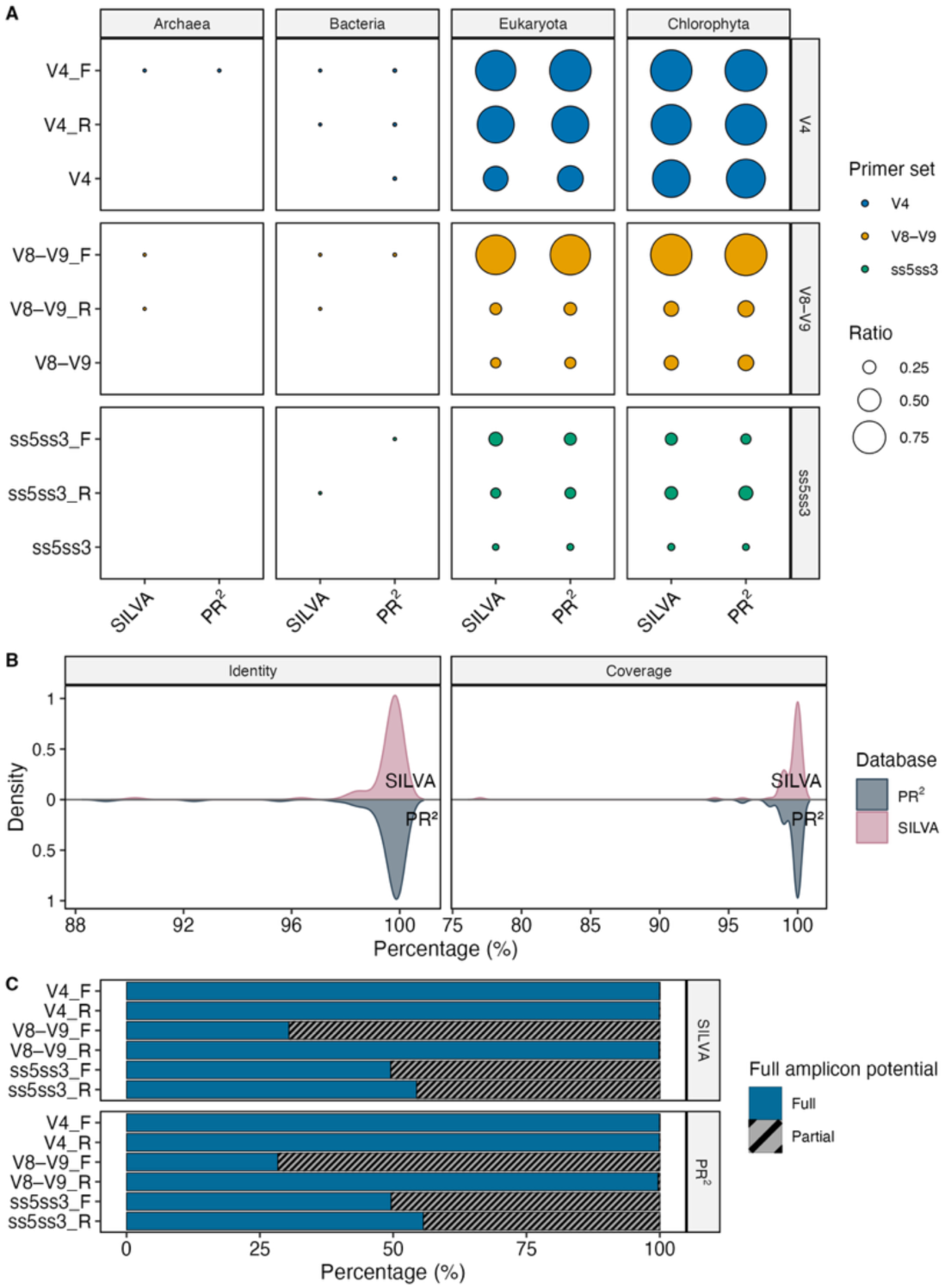
In silico primer coverage analysis across SILVA and PR² databases. (A) Proportional coverage of each primer set (V4, blue; V8–V9, orange; ss5ss3, green) for main taxonomic groups (Archaea, Bacteria, Eukaryota, and Chlorophyta); bubble size indicates the coverage ratio. (B) Identity and query coverage distributions of BLAST best hits for PR² bacterial sequences amplified by the V4 primer, shown as mirrored density curves; SILVA hits (pink, upper) and PR² eukaryote hits (gray, lower). (C) Full amplicon potential of individual primers across SILVA and PR² databases. Bars represent the percentage of single-primer-matched sequences sufficiently long to accommodate the full amplicon; blue bars (solid) indicate sequences with full amplicon potential and grey bars (hatched) indicate sequences without full amplicon potential.

Primer pair coverage was assessed across major taxonomic groups (Bacteria, Archaea, Eukaryota, and Chlorophyta). The V4 primer pair, consistent with its superior performance in the initial coverage analysis, was observed to exhibit the highest taxonomic coverage among eukaryotic sequences in both reference databases (SILVA: 54.94%, PR^2^: 57.71%). This trend was notably pronounced for Chlorophyta sequences (SILVA: 87.76%, PR^2^: 91.25%), the principal group in green algae. In contrast, V8–V9 and ss5ss3 primer pairs were found to demonstrate substantially lower coverage, with only 17.63% and 20.11% of eukaryotic sequences, and 28.18% and 31.54% of Chlorophyta sequences in SILVA and PR^2^, respectively. The ss5ss3 pair exhibited even lower coverage, capturing only 8.38% and 9.45% of eukaryotic sequences, and 10.15% and 9.11% of Chlorophyta sequences. Further analysis of taxonomic specificity revealed that the combined primer pairs were generally specific to eukaryotic 18S rRNA genes. V8–V9 and ss5ss3 primer pairs did not capture any 16S rRNA gene sequences, while the V4 primer pair captured 78 sequences annotated as bacterial sequences in the PR^2^ database (Figure 2A). However, these sequences exhibited high-identity matches to eukaryotic sequences in both databases (SILVA: 99.52 ± 1.21%, PR^2^: 99.43 ± 1.58%) with extensive query coverage (SILVA: 99.33 ± 2.71%, PR^2^: 99.54 ± 1.03%) (Figure 2B). These findings suggest taxonomic mis-annotations in the published reference databases.

*In silico* analysis has become a widely adopted tool for initial assessments of primer coverage and specificity in microbiome research.^36,37,68,69^ While these computational approaches offer valuable preliminary insights, our study reveals significant limitations in publicly available reference databases that warrant careful consideration. Our investigation identified two critical issues in these databases. First, the presence of partial 18S rRNA gene sequences can significantly impact primer selection and study design. For instance, primers targeting terminal regions or spanning nearly the entire gene, such as V8–V9 and ss5ss3, may appear less effective *in silico* than they perform *in vitro* due to incomplete reference sequences. This bias can lead to underestimation of primer utility in computational analyses. Second, we discovered instances of taxonomic mis-annotations in the PR^2^ database, specifically eukaryotic sequences being incorrectly labeled as bacteria. This finding is particularly concerning as it can lead to potential distortions in our understanding of microbial community composition and diversity.^70^ Previous studies have reported mis-annotation rates ranging from 0.2% to 20% across various microbial reference databases,^71–74^ indicating that this is a widespread issue in the field. These limitations underscore the need for cautious interpretation of all results dependent on reference databases.

### 18S rRNA gene analysis indicated primer pair and amplicon length dependent insights into eukaryotic community

Sequencing of samples from the EcoRecover process resulted in 9,146,557 SR sequences and 1,444,452 LR sequences from 48 and 47 samples, producing 2,369 and 2,799 ASVs, respectively. *In silico* PCR analysis identified V8–V9 amplicons in 2,798 of 2,799 LR ASVs, which were subsequently de-replicated to 462 ASVs. After technical duplicate merging, filtering, and rarefaction, the final datasets contained 2,845,380 sequences (1,091 ASVs), 326,790 sequences (1,150 ASVs), and 339,180 sequences (241 ASVs) from 30 samples from the SR, LR, and LS datasets, respectively. The reduction from the initial 48 samples to the final 30 samples occurred during the rarefaction process, where samples with insufficient sequencing depth were excluded. After rarefaction, only 30 samples remained common across all datasets, allowing for direct comparative analyses. SR and LR exhibited comparable richness (310.27 ± 60.75 and 359.57 ± 175.87 ASVs), while LS showed significantly lower richness (55.13 ± 35.31 ASVs) (Figure S1A). LR demonstrated the highest Shannon diversity (4.19 ± 0.66) and Simpson index (0.91 ± 0.10), followed by SR (2.34 ± 0.47; 0.74 ± 0.10) and LS (1.57 ± 0.41; 0.62 ± 0.14) (Figure 3A, S1B). Statistical analysis revealed significant differences between sequencing approaches (p < 0.001), except for the SR versus LR comparison in observed ASVs.

**Figure 3:**
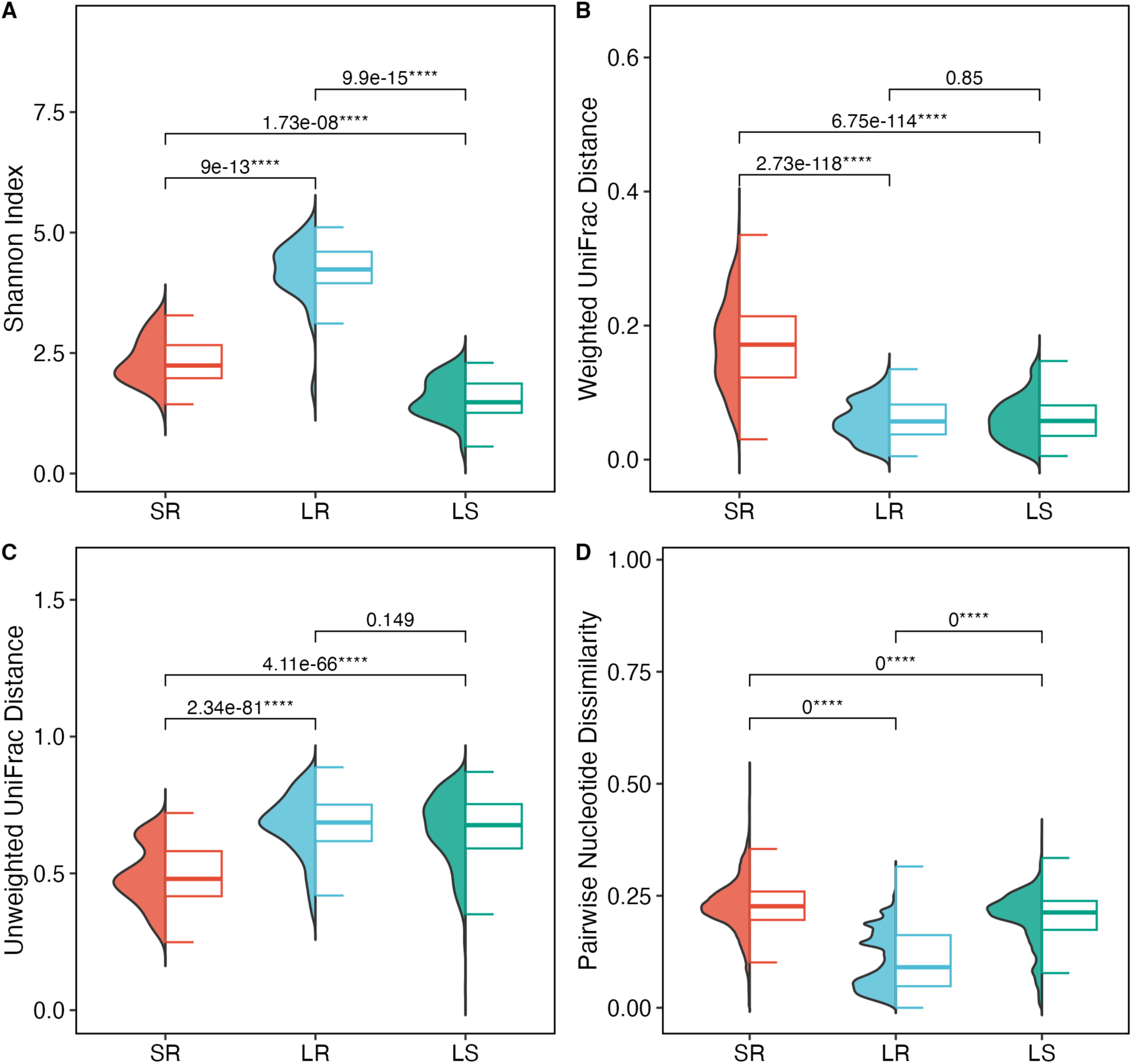
Comparison of diversity metrics across three 18S rRNA gene sequencing datasets: (A) Shannon diversity index; (B) weighted UniFrac distances; (C) unweighted UniFrac distances; (D) pairwise ASV nucleotide dissimilarities. SR (red): short-read V8–V9 region sequenced using Illumina platform; LR (blue): long-read full-length 18S rRNA gene sequenced using PacBio platform; LS (green): computationally extracted V8–V9 regions from the LR dataset. Half-violin plots illustrate the density distribution of each metric; half-boxplots show median and interquartile range. Pairwise differences were assessed using Wilcoxon tests with Benjamini-Hochberg correction.

Analysis of computationally extracted V8–V9 regions (i.e., LS dataset) from the LR dataset showed that V8–V9 primers matched 99.96% of ss5ss3 amplicon sequences (2,798 out of 2,799). While this indicates high theoretical amplification efficiency of V8–V9 primers for ss5ss3-targeted populations, sequence dereplication yielded only 462 unique ASVs. The decreased richness observed in the V8–V9 region (∼375 bp) reflects its reduced taxonomic resolution, as multiple species can share identical sequences within this shorter fragment. The LR dataset exhibited comparable richness levels to the SR dataset, despite ss5ss3 primers targeting a narrower taxonomic range. The observed diversity pattern is likely due to increased sequence variation captured across the near-full-length 18S rRNA gene (∼1800 bp). While LR and SR demonstrated comparable richness, their diversity measures using Shannon and Simpson indices differed substantially. The higher diversity in LR datasets indicates that the community captured by the ss5ss3 primer pair and sequenced using long-read methods displays higher evenness, likely resulting from the enhanced taxonomic resolution of the near-full-length 18S rRNA gene providing better discrimination of closely related taxa. Beta-diversity analyses using multiple dissimilarity metrics revealed distinct patterns among sequencing approaches. Bray-Curtis dissimilarity analysis showed that LR samples exhibited the highest average dissimilarity (0.62 ± 0.22), followed by SR (0.53 ± 0.22) and LS samples (0.45 ± 0.23), with statistically significant differences across all approaches (p < 1e^-4^) (**Figure S2**). When incorporating phylogenetic information through UniFrac distances, both weighted and unweighted UniFrac analyses demonstrated high consistency between LR and LS datasets. The weighted UniFrac dissimilarity values were nearly identical for LR (0.059 ± 0.029) and LS (0.060 ± 0.032), while SR showed significantly higher dissimilarity (0.171 ± 0.063, p < 1e^-4^) (**Figure 3B**). Similarly, unweighted UniFrac analysis revealed comparable values between LR (0.68 ± 0.11) and LS (0.66 ± 0.13), though in this case, SR exhibited significantly lower dissimilarity (0.49 ± 0.11, p < 1e^-4^) (Figure 3C). The comparable dissimilarity between LR and LS using both weighted and unweighted UniFrac distances indicates phylogenetic relationships are preserved when extracting shorter regions from full-length sequences, while the contrasting comparison to SR datasets indicates V8–V9 primers capture phylogenetically distinct abundant taxa but more similar rare taxa.

The findings based on alpha and beta-diversity analysis were further supported by pairwise ASV dissimilarity (Figure 3D). LR dataset showed the lowest dissimilarity (0.11 ± 0.07), followed by LS (0.20 ± 0.06) and SR (0.23 ± 0.06) datasets, with significant differences among all three approaches (p < 1e^-4^). The identity distribution patterns revealed distinct characteristics: LR dataset exhibited multiple distinct peaks, indicating several phylogenetic clusters with closely related ASVs but limited between-cluster similarity. Despite LS being derived from LR sequences, the identity distribution pattern of the LS dataset more closely resembled the SR dataset, suggesting that the V8–V9 region alone had a reduced taxonomic resolution compared to full-length sequences.

The relationships between different sequencing approaches were further evaluated using Mantel tests based on both weighted and unweighted UniFrac distances (Figure 4). All pairwise comparisons demonstrated significant correlations (p = 0.001), indicating that the community structures detected by different sequencing approaches were correlated. However, correlation coefficients were moderate (R = 0.423–0.607) for comparisons involving SR, suggesting substantial variation in community structure detection across sequencing approaches. The strongest correlation was observed between LR and LS datasets (weighted UniFrac: R = 0.933; unweighted UniFrac: R = 0.804), which is expected since the LS dataset was computationally derived from the LR dataset. The consistently lower correlation coefficients in unweighted UniFrac comparisons (SR-LR: R = 0.561; SR-LS: R = 0.423) compared to their weighted counterparts (SR-LR: R = 0.607; SR-LS: R = 0.456) suggest that abundance-weighted community structures are more consistent across different sequencing approaches. These moderate yet significant correlations indicate that while these methods capture similar broad community patterns, there are notable methodological biases that should be considered when selecting sequencing approaches for microalgal community analysis.

**Figure 4:**
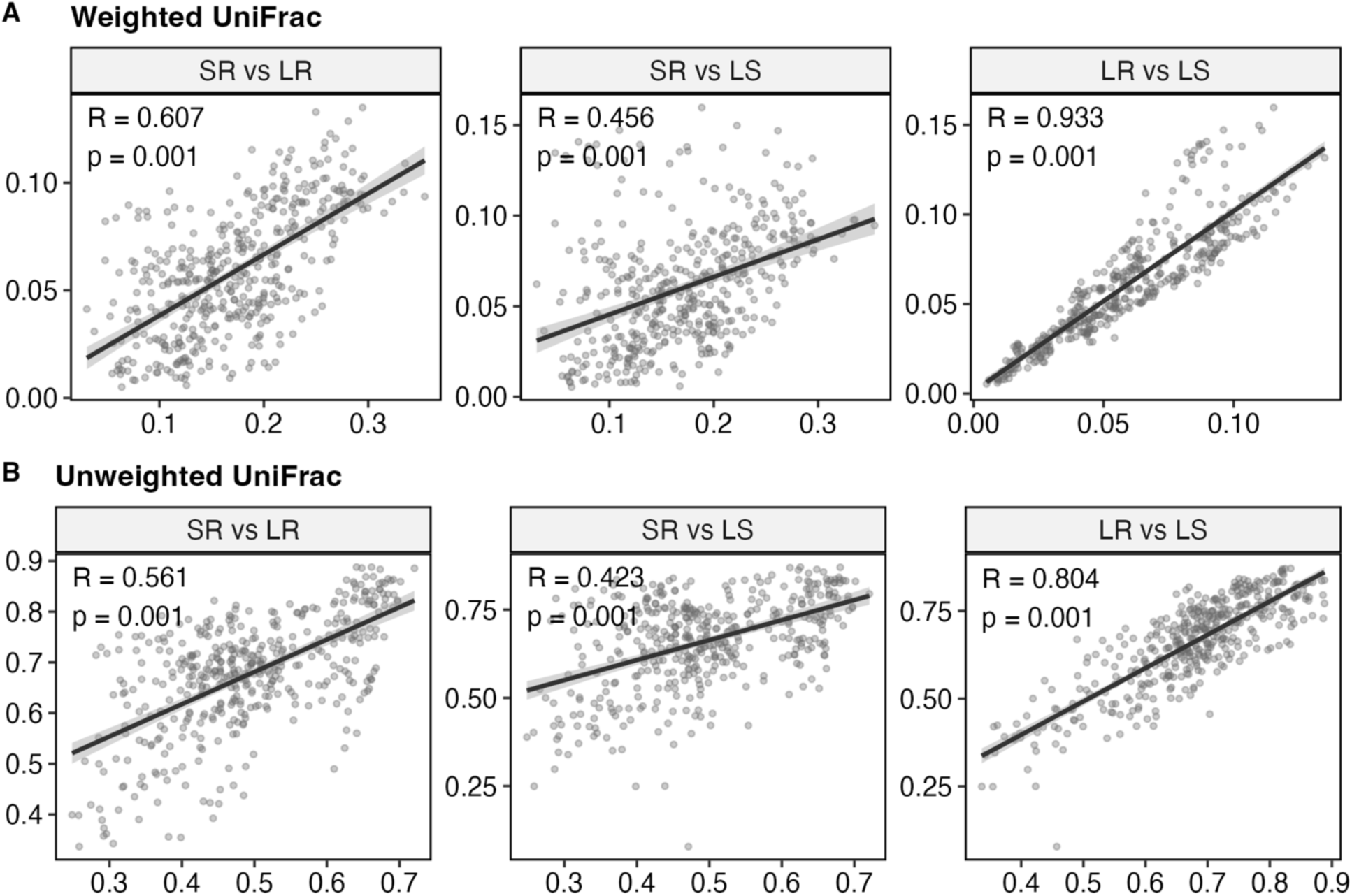
Correlation analysis of UniFrac distances between sequencing approaches. (A) Weighted UniFrac distances. (B) Unweighted UniFrac distances. Lines represent linear regression fits with 95% confidence intervals (shaded). R and p-values are from Spearman Mantel tests (999 permutations). SR: short-read V8–V9 region sequenced using Illumina platform; LR: long-read full-length 18S rRNA gene sequenced using PacBio platform; LS: computationally extracted V8–V9 regions from the LR dataset.

### Amplicon classification outcomes is database dependent

Using the SILVA database with QIIME2’s Naive Bayes classifier, LR maintained high classified count percentage and relative abundance through the class level (98.5%; 99.5% ± 0.51%), whereas SR and LS declined markedly at the phylum level in both classified count percentage (80.3% and 78.4%, respectively) and relative abundance (84.2% ± 23.1% and 78.1% ± 25.8%, respectively). However, LR showed no advantage at higher taxonomic resolution with SILVA: below the order level, LR relative abundance fell below that of SR (order: 68.7% vs. 80.1%), and at the genus and species levels, LR also exhibited lower classified count percentages than SR (genus: 42.6% vs. 51.6%; species: 33.2% vs. 35.6%) (Figure 5, Table S1). This observation may be attributed to two factors. First, Chlorophyta, the predominant group in EcoRecover samples, accounts for only 0.37% of SILVA sequences (3.25% of eukaryotic sequences), which may limit classification of this key group. Second, SILVA’s curation extends only to the genus level, and approximately 72% of species labels have been reported as unidentified or uncultured organisms, with an additional 2.5% exhibiting genus–species mismatches, reducing classification accuracy below the genus level.^46^ To address these limitations, the RDP classifier with the PR^2^ database was implemented. PR^2^ is a eukaryote-focused database with curated species-level annotation, in which Chlorophyta comprises 5.40% of total sequences, directly benefiting LR’s full-length 18S rRNA gene sequences that require species-level reference matches. With PR^2^, LR outperformed both SR and LS across all taxonomic ranks. At the domain level, all methods achieved 100% classified count percentage. From supergroup to class, LR maintained high classified count percentages (98.5% to 93.7%), whereas SR declined markedly (88.7% to 64.2%) and LS showed intermediate performance (94.6% to 61.0%). At the genus and species levels, LR maintained substantially higher relative abundances than SR and LS (genus: 80.8% vs. 32.6% vs. 18.3%; species: 65.4% vs. 13.1% vs. 4.5%) (Figure 5A, Table S1). These results indicate that realizing the full potential of long-read sequencing requires appropriate reference database selection. PR^2^’s species-level curation and comprehensive Chlorophyta representation enable taxonomic assignment of full-length 18S rRNA gene sequences, improving classified count percentages and microalgal community characterization across all taxonomic ranks.

**Figure 5:**
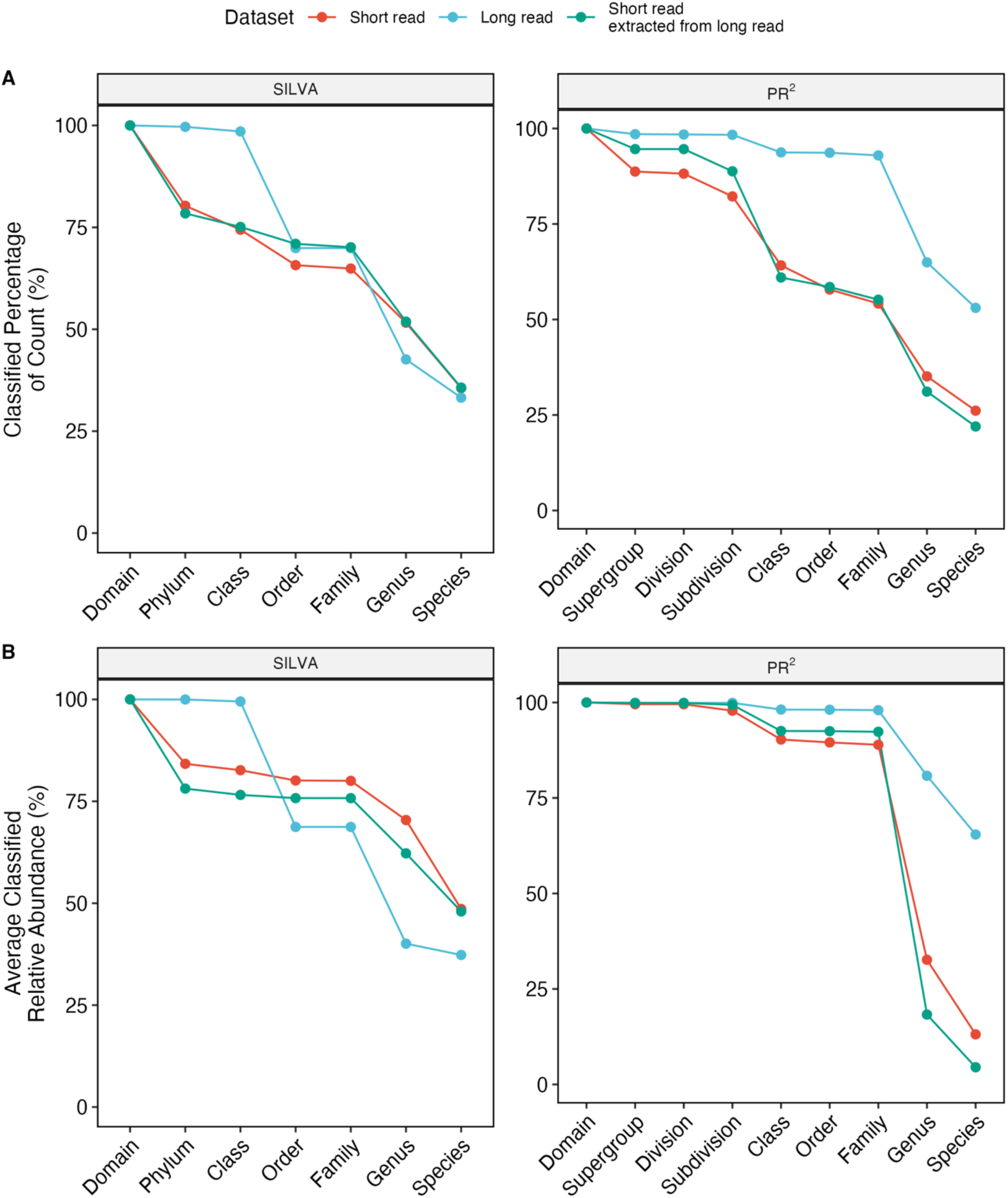
Performance comparison of taxonomic classification approaches across sequencing datasets. (A) Percentage of classified sequences and (B) relative abundance of classified taxa at each taxonomic rank, using QIIME2’s Naive Bayes classifier with SILVA database and RDP classifier with PR² database. SR (red): short-read V8–V9 region sequenced using Illumina platform; LR (blue): long-read full-length 18S rRNA gene sequenced using PacBio platform; LS (green): computationally extracted V8–V9 regions from the LR dataset.

### Impact of Sequencing Approaches on Process Relevant Insights

The Roberts wastewater treatment plant (WWTP)’s Wisconsin Pollutant Discharge Elimination System (WPDES) permit (No. 0028835) requires effluent orthophosphate (OP) ≤0.04 mg-P·L⁻¹, down from 1 mg-P·L⁻¹ prior to February 1, 2021.^6^ Samples from the EcoRecover process were classified as “Achieved” (Y; ≤0.04 mg-P·L^-1^) or “Not Achieved” (N; >0.04 mg-P·L^-1^) based on effluent OP concentration (Figure S3). Of the 30 samples analyzed, 16 achieved the target limit while 14 did not meet the requirement. Time series analysis revealed significant temporal variations in effluent orthophosphate concentrations during the monitoring period (August 2022 to August 2023), with concentrations ranging from 0 to 3.57 mg-P·L^-1^ (mean: 0.43 ± 0.78 mg-P·L^-1^).

Alpha diversity metrics were compared between Y and N groups across three sequencing datasets (Figure 6A). All three approaches (SR, LR, and LS) showed comparable results for Shannon and Simpson indices, with no significant differences observed between Y and N groups (p > 0.05). Beta diversity analysis using weighted and unweighted UniFrac distances revealed complex community structuring patterns (Figure 6B). Principal coordinate analysis (PCoA) demonstrated substantial variation explained by the first two axes in both weighted (40.23–61.70%) and unweighted (36.51–42.72%) UniFrac analyses. While visual inspection of PCoA plots showed no clear group segregation, permutational multivariate analysis of variance (PERMANOVA) analysis uncovered subtle differences. The unweighted UniFrac analysis detected weak but significant differences between Y and N groups in LR (R^2^ = 0.074, p = 0.015) and LS (R^2^ = 0.075, p = 0.03) datasets, with these differences explaining less than 8% of community variation. In contrast, weighted UniFrac analysis showed no significant differences across all datasets (SR: R^2^ = 0.018, p = 0.756; LR: R^2^ = 0.03, p = 0.434; LS: R^2^ = 0.054, p = 0.172). These results suggest that while rare taxa might exhibit subtle variations between groups in LR and LS datasets, the abundant taxa and overall community structure remained largely consistent. The consistency between Y and N groups across all three sequencing datasets indicates that while full-length sequencing provides enhanced taxonomic resolution, the selection of sequencing method does not influence operational performance assessment. These results suggest that shorter amplicon sequencing would be adequate for routine operational monitoring, offering a cost-effective approach despite its lower taxonomic resolution.

**Figure 6:**
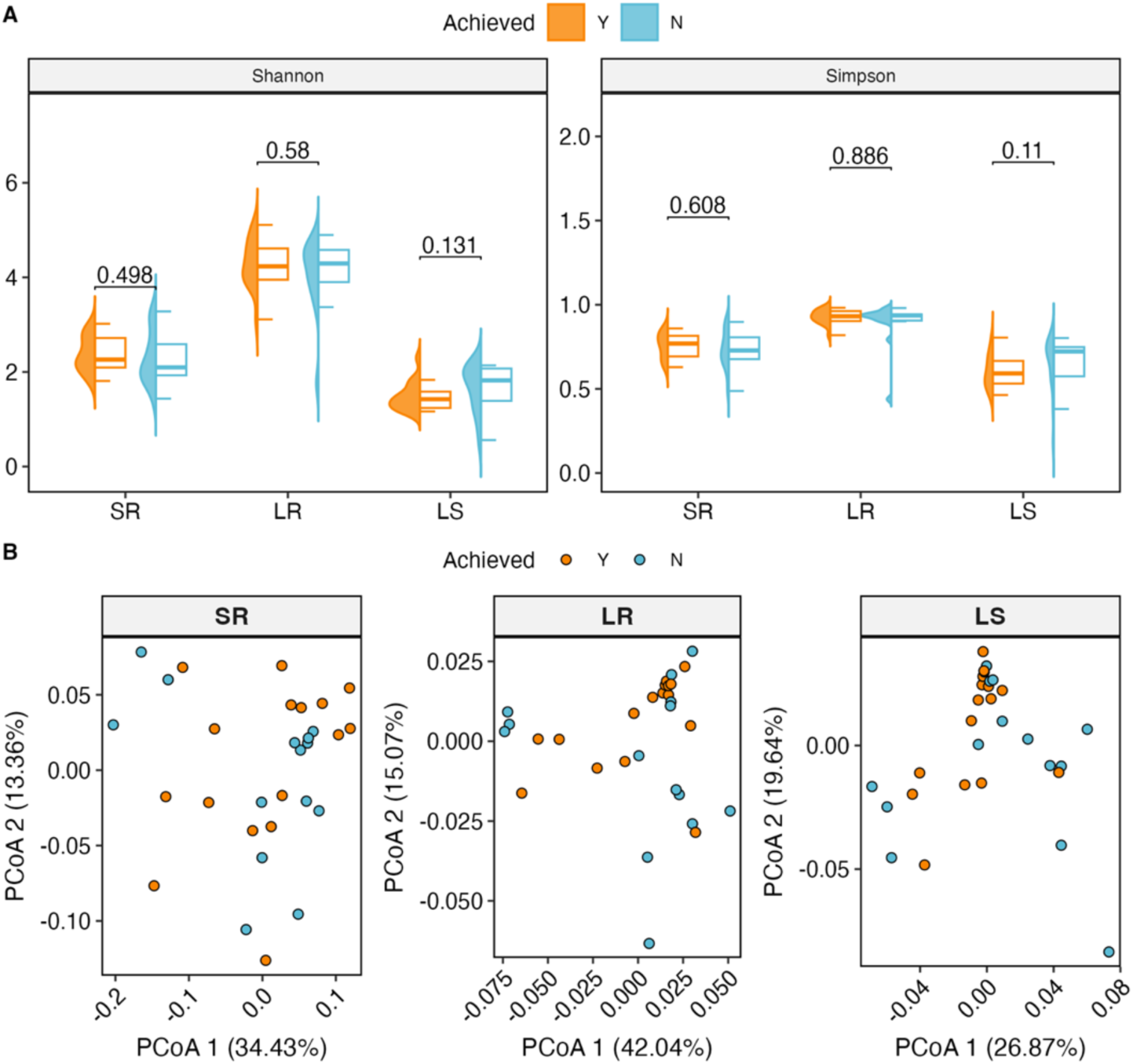
Alpha and beta diversity analyses of microalgal communities across three sequencing datasets, grouped by phosphorus removal performance (Y: achieved ≤ 0.04 mg-P·L⁻¹; N: not achieved). (A) Half-violin and half-boxplot comparison of Shannon and Simpson diversity indices. P-values from Wilcoxon tests with Benjamini-Hochberg correction. (B) Principal Coordinate Analysis (PCoA) based on weighted UniFrac distances for SR (short-read V8–V9, Illumina), LR (full-length 18S, PacBio), and LS (V8–V9 extracted from LR) datasets. Orange symbols: Y group; blue symbols: N group.

### Impact of sequencing approaches on differential abundance analysis

Samples from two specific periods (August 10, 2022 – November 2, 2022 and December 8, 2022 – April 28, 2023) were selected for differential abundance analyses to identify ASVs associated with OP removal performance, as substantial temporal variation across the full dataset would otherwise obscure underlying patterns in microalgal community structure. Full-length 18S rRNA gene sequencing (LR) provided substantially greater taxonomic resolution within Chlorophyta than either the in vitro V8–V9 amplicon dataset (SR) or the *in silico*-extracted V8–V9 region (LS). All 15 LR ASVs enriched in Y group samples were resolved to species level, identified as *Scenedesmus obliquus* (n = 9), *Desmodesmus pannonicus* (n = 3), and *Acutodesmus* sp. (n = 3). Both short-read approaches performed markedly worse: the SR dataset assigned all 7 Chlorophyta ASVs enriched in Y group samples only to family or subdivision level (Sphaeropleales_X, n = 2; Chlorellales_X, n = 3; Chlorophyta_X, n = 2), and the LS dataset resolved only 1 of 8 corresponding ASVs to species level (Desmodesmus opoliensis), with the remaining ASVs classified only to family (Sphaeropleales_X, n = 4) or subdivision level (Chlorophyta_X, n = 3). A similar pattern was observed in N group samples: the LR dataset resolved 4 of 6 ASVs to genus level (*Dictyosphaerium* sp., n = 3; *Dictyosphaerium*, n = 1), with 2 remaining at family level (Chlorellales_X), while both LS ASVs and the SR Chlorophyta ASVs enriched in N group were classified only to family level (Chlorellales_X) (Tables S2, S3).

*Scenedesmus*, *Desmodesmus*, and *Acutodesmus* are well-documented phosphorus assimilators across diverse wastewater streams, including piggery wastewater, dairy effluent, aquaculture wastewater, tannery effluent, and municipal wastewater.^75–81^ *Desmodesmus* sp. has demonstrated efficient nitrogen and phosphorus removal in photobioreactor-based wastewater treatment, with phosphate taken up in excess of immediate cellular requirements, indicative of luxury uptake.^82^ *Scenedesmus obliquus* similarly exhibits dynamic phosphorus uptake in photobioreactor systems, with carbohydrate mobilization sustaining assimilation during dark periods.^6,83^ Despite belonging to the same family (Sphaeropleales), *Desmodesmus* sp. and Scenedesmus obliquus exhibit distinct phosphorus removal capacities, and family-level classification alone, as produced by both SR and LS, would collapse them into an unresolved unit (Sphaeropleales_X), obscuring the underlying removal mechanisms.

The SR dataset, which targeted the V8–V9 region, extended coverage beyond Chlorophyta to include heterotrophic taxa not captured by LR or LS sequencing. ASVs assigned to Bicoecaceae and unresolved fungal organisms were enriched in both Y and N groups, represented by distinct ASVs in each. An ASV assigned to *Spumella* was exclusively enriched in Y group samples, while a separate ASV resolved to *Spumella vulgaris* was exclusively enriched in N group samples. A single ASV assigned to *Eucyclops serrulatus* was enriched in Y group samples. ASVs assigned to Vampyrellida (Leptophryidae), Cryomonadida (Rhogostoma lineage), and Choanoflagellata (Craspedida, including one ASV resolved to *Monosiga ovata*) were exclusively enriched in N group samples (Tables S3, S4).

Members of Leptophryidae are documented contaminants of microalgal cultivation systems. *Vernalophrys algivore*, isolated from outdoor *Scenedesmus dimorphous* mass cultures, was reported as a significant grazer associated with rapid algal cell number declines,^84^ and *Kinopus chlorellivorus* was identified as a direct cause of biomass loss in large-scale Chlorella cultures, with the capacity to consume *Scenedesmus* and other chlorophyte microalgae.^85^ The enrichment of this ASV in N group samples therefore suggests a potential role in suppressing the Sphaeropleales-dominated community responsible for phosphorus removal. ASVs assigned to Choanoflagellata (Craspedida) and the Rhogostoma lineage were also exclusively enriched in N group samples. These taxa are known bacterial predators.^86–88^ The association of Rhogostoma-lineage ASVs with reduced performance is further supported by prior observations from the same facility, in which Cercozoa of this lineage proliferated ahead of rotifer blooms during periods of declining system performance.^12^ Together, the enrichment of these heterotrophic bacterivores suggest a community shift away from phototrophic dominance, with elevated bacterial prey availability potentially associated with reduced algal biomass. LR sequencing thus provided the species-level resolution necessary to identify the algal taxa functionally responsible for phosphorus removal, while SR sequencing revealed a broader assemblage of heterotrophic predators and bacterivores whose enrichment was associated with reduced performance.

## Conclusion

Short-read sequencing using the V8–V9 primers provided broader taxonomic coverage and effectively captured the community members essential for system performance assessment. This approach also captured algal-predatory and bacterivorous protists enriched when performance declined, suggesting grazing pressure on the phosphorus-removing community. Long-read sequencing achieved superior taxonomic classification resolution, enabling species-level identification of functionally distinct taxa within Chlorophyta which would remain unresolved by short-read approaches alone. Method selection should therefore be guided by research objective: short-read sequencing for broader community profiling across diverse eukaryotic groups, and long-read sequencing for species-level identification of functional taxa. This study also highlights critical limitations in current reference databases, including partial reference sequences and taxonomic mis-annotations. These deficiencies compromise *in silico* performance assessments and taxonomic classification accuracy, preventing long-read sequencing from fully realizing its taxonomic resolution potential. Future improvements in reference database quality and curation practices will enhance the utility of both sequencing approaches for environmental engineering applications. These findings provide practical guidance for researchers and operators selecting appropriate sequencing methods for monitoring and optimizing microalgal-based wastewater treatment systems.

## Author Contributions

**Zihan Dai**: writing−original draft, writing−review and editing, formal analysis, data curation, methodology, investigation, and conceptualization. **Md Mahbubul Alam**: resources and writing−review and editing. **Benjamin Gincley**: resources and writing−review and editing. **Farhan Khan**: resources and writing−review and editing. **Ga-Yeong Kim**: resources and writing−review and editing. **Hannah Molitor**: resources and writing−review and editing. **Jeremy Guest**: supervision, funding acquisition, and writing−review and editing. **Ian M. Bradley**: supervision, funding acquisition, and writing−review and editing. **Ameet J. Pinto**: supervision, funding acquisition, conceptualization, and writing−review and editing.

## Acknowledgements

This work was supported by the U.S. Department of Energy’s Office of Energy Efficiency and Renewable Energy (EERE) Bioenergy Technologies Office (BETO) under the FY20 Bioenergy Technologies Multi-Topic FOA Award Number(s) DE-EE0009270. Claude (Anthropic, claude.ai) was used to assist with improving the flow and readability of the manuscript text and with R script formatting, annotation, and reproducibility improvements. All scientific content, data interpretation, and conclusions were developed solely by the authors. The authors take full responsibility for all submitted content.

## References

(1) Cai, T.; Park, S. Y.; Li, Y. Nutrient Recovery from Wastewater Streams by Microalgae: Status and Prospects. Renew. Sustain. Energy Rev. 2013, 19, 360–369. 10.1016/j.rser.2012.11.030.

(2) Gonçalves, A. L.; Pires, J. C. M.; Simões, M. A Review on the Use of Microalgal Consortia for Wastewater Treatment. Algal Res. 2017, 24, 403–415. 10.1016/j.algal.2016.11.008.

(3) Park, J. B. K.; Craggs, R. J.; Shilton, A. N. Wastewater Treatment High Rate Algal Ponds for Biofuel Production. Bioresour. Technol. 2011, 102 (1), 35–42. 10.1016/j.biortech.2010.06.158.

(4) Bora, A.; Thondi Rajan, A. S.; Ponnuchamy, K.; Muthusamy, G.; Alagarsamy, A. Microalgae to Bioenergy Production: Recent Advances, Influencing Parameters, Utilization of Wastewater – a Critical Review. Sci. Total Environ. 2024, 946, 174230. 10.1016/j.scitotenv.2024.174230.

(5) Mohsenpour, S. F.; Hennige, S.; Willoughby, N.; Adeloye, A.; Gutierrez, T. Integrating Micro-Algae into Wastewater Treatment: A Review. Sci. Total Environ. 2021, 752, 142168. 10.1016/j.scitotenv.2020.142168.

(6) Molitor, H. R.; Kim, G.-Y.; Hartnett, E.; Gincley, B.; Alam, M. M.; Feng, J.; Avila, N. M.; Fisher, A.; Hodaei, M.; Li, Y.; McGraw, K.; Cusick, R. D.; Bradley, I. M.; Pinto, A. J.; Guest, J. S. Intensive Microalgal Cultivation and Tertiary Phosphorus Recovery from Wastewaters via the EcoRecover Process. Environ. Sci. Technol. 2024, 58 (20), 8803–8814. 10.1021/acs.est.3c10264.

(7) Abdelfattah, A.; Ali, S. S.; Ramadan, H.; El-Aswar, E. I.; Eltawab, R.; Ho, S.-H.; Elsamahy, T.; Li, S.; El-Sheekh, M. M.; Schagerl, M.; Kornaros, M.; Sun, J. Microalgae-Based Wastewater Treatment: Mechanisms, Challenges, Recent Advances, and Future Prospects. Environ. Sci. Ecotechnol. 2023, 13, 100205. 10.1016/j.ese.2022.100205.

(8) Amaro, H. M.; Salgado, E. M.; Nunes, O. C.; Pires, J. C. M.; Esteves, A. F. Microalgae Systems - Environmental Agents for Wastewater Treatment and Further Potential Biomass Valorisation. J. Environ. Manage. 2023, 337, 117678. 10.1016/j.jenvman.2023.117678.

(9) Zhou, Q.; Sun, H.; Jia, L.; Wu, W.; Wang, J. Simultaneous Biological Removal of Nitrogen and Phosphorus from Secondary Effluent of Wastewater Treatment Plants by Advanced Treatment: A Review. Chemosphere 2022, 296, 134054. 10.1016/j.chemosphere.2022.134054.

(10) Nguyen, L. N.; Aditya, L.; Vu, H. P.; Johir, A. H.; Bennar, L.; Ralph, P.; Hoang, N. B.; Zdarta, J.; Nghiem, L. D. Nutrient Removal by Algae-Based Wastewater Treatment. Curr. Pollut. Rep. 2022, 8 (4), 369–383. 10.1007/s40726-022-00230-x.

(11) Razzak, S. A. Recent Advances in Sustainable Biological Nutrient Removal from Municipal Wastewater. *Clean*. Water 2024, 2, 100047. 10.1016/j.clwat.2024.100047.

(12) Alam, M. M.; Hodaei, M.; Hartnett, E.; Gincley, B.; Khan, F.; Kim, G.-Y.; Pinto, A. J.; Bradley, I. M. Community Structure and Function during Periods of High Performance and System Upset in a Full-Scale Mixed Microalgal Wastewater Resource Recovery Facility. Water Res. 2024, 259, 121819. 10.1016/j.watres.2024.121819.

(13) Oruganti, R. K.; Katam, K.; Show, P. L.; Gadhamshetty, V.; Upadhyayula, V. K. K.; Bhattacharyya, D. A Comprehensive Review on the Use of Algal-Bacterial Systems for Wastewater Treatment with Emphasis on Nutrient and Micropollutant Removal. Bioengineered 2022, 13 (4), 10412–10453. 10.1080/21655979.2022.2056823.

(14) Mattsson, L.; Sörenson, E.; Capo, E.; Farnelid, H. M.; Hirwa, M.; Olofsson, M.; Svensson, F.; Lindehoff, E.; Legrand, C. Functional Diversity Facilitates Stability under Environmental Changes in an Outdoor Microalgal Cultivation System. Front. Bioeng. Biotechnol. 2021, 9, 651895. 10.3389/fbioe.2021.651895.

(15) Novoveská, L.; Zapata, A. K. M.; Zabolotney, J. B.; Atwood, M. C.; Sundstrom, E. R. Optimizing Microalgae Cultivation and Wastewater Treatment in Large-Scale Offshore Photobioreactors. Algal Res. 2016, 18, 86–94. 10.1016/j.algal.2016.05.033.

(16) Liu, J.; Wu, Y.; Wu, C.; Muylaert, K.; Vyverman, W.; Yu, H.-Q.; Muñoz, R.; Rittmann, B. Advanced Nutrient Removal from Surface Water by a Consortium of Attached Microalgae and Bacteria: A Review. Bioresour. Technol. 2017, 241, 1127–1137. 10.1016/j.biortech.2017.06.054.

(17) De Morais, E. G.; Sampaio, I. C. F.; Gonzalez-Flo, E.; Ferrer, I.; Uggetti, E.; García, J. Microalgae Harvesting for Wastewater Treatment and Resources Recovery: A Review. New Biotechnol. 2023, 78, 84–94. 10.1016/j.nbt.2023.10.002.

(18) Benedetti, M.; Vecchi, V.; Barera, S.; Dall’Osto, L. Biomass from Microalgae: The Potential of Domestication towards Sustainable Biofactories. Microb. Cell Fact. 2018, 17 (1), 173–190. 10.1186/s12934-018-1019-3.

(19) Sutherland, D. L.; Howard-Williams, C.; Turnbull, M. H.; Broady, P. A.; Craggs, R. J. The Effects of CO2 Addition along a pH Gradient on Wastewater Microalgal Photo-Physiology, Biomass Production and Nutrient Removal. Water Res. 2015, 70, 9–26. 10.1016/j.watres.2014.10.064.

(20) Krustok, I.; Odlare, M.; Truu, J.; Nehrenheim, E. Inhibition of Nitrification in Municipal Wastewater-Treating Photobioreactors: Effect on Algal Growth and Nutrient Uptake. Bioresour. Technol. 2016, 202, 238–243. 10.1016/j.biortech.2015.12.020.

(21) Bukin, Y. S.; Mikhailov, I. S.; Petrova, D. P.; Galachyants, Y. P.; Zakharova, Y. R.; Likhoshway, Y. V. The Effect of Metabarcoding 18S rRNA Region Choice on Diversity of Microeukaryotes Including Phytoplankton. World J. Microbiol. Biotechnol. 2023, 39 (9), 229–245. 10.1007/s11274-023-03678-1.

(22) Bradley, I. M.; Pinto, A. J.; Guest, J. S. Design and Evaluation of Illumina MiSeq-Compatible, 18S rRNA Gene-Specific Primers for Improved Characterization of Mixed Phototrophic Communities. Appl. Environ. Microbiol. 2016, 82 (19), 5878–5891. 10.1128/AEM.01630-16.

(23) Gaonkar, C. C.; Campbell, L. Metabarcoding Reveals High Genetic Diversity of Harmful Algae in the Coastal Waters of Texas, Gulf of Mexico. Harmful Algae 2023, 121, 102368. 10.1016/j.hal.2022.102368.

(24) Ferro, L.; Hu, Y. O. O.; Gentili, F. G.; Andersson, A. F.; Funk, C. DNA Metabarcoding Reveals Microbial Community Dynamics in a Microalgae-Based Municipal Wastewater Treatment Open Photobioreactor. Algal Res. 2020, 51, 102043. 10.1016/j.algal.2020.102043.

(25) Remias, D.; Procházková, L.; Nedbalová, L.; Benning, L. G.; Lutz, S. Novel Insights in Cryptic Diversity of Snow and Glacier Ice Algae Communities Combining 18S rRNA Gene and ITS2 Amplicon Sequencing. FEMS Microbiol. Ecol. 2023, 99 (12), fiad134. 10.1093/femsec/fiad134.

(26) Olefeld, J. L.; Bock, C.; Jensen, M.; Vogt, J. C.; Sieber, G.; Albach, D.; Boenigk, J. Centers of Endemism of Freshwater Protists Deviate from Pattern of Taxon Richness on a Continental Scale. Sci. Rep. 2020, 10 (1), 14431. 10.1038/s41598-020-71332-z.

(27) Jacobs-Palmer, E.; Gallego, R.; Cribari, K.; Keller, A. G.; Kelly, R. P. Environmental DNA Metabarcoding for Simultaneous Monitoring and Ecological Assessment of Many Harmful Algae. Front. Ecol. Evol. 2021, 9, 612107. 10.3389/fevo.2021.612107.

(28) Kravtsova, L. S.; Peretolchina, T. E.; Triboy, T. I.; Nebesnykh, I. A.; Kupchinskiy, A. B.; Tupikin, A. E.; Kabilov, M. R. The Study of the Diversity of Hydrobionts from Listvennichny Bay of Lake Baikal by DNA Metabarcoding. Russ. J. Genet. 2021, 57 (4), 460–467. 10.1134/S1022795421040050.

(29) Vieira, H. H.; Bagatini, I. L.; Guinart, C. M.; Vieira, A. A. H. tufA Gene as Molecular Marker for Freshwater Chlorophyceae. ALGAE 2016, 31 (2), 155–165. 10.4490/algae.2016.31.4.14.

(30) Ballesteros, I.; Terán, P.; Guamán-Burneo, C.; González, N.; Cruz, A.; Castillejo, P. DNA Barcoding Approach to Characterize Microalgae Isolated from Freshwater Systems in Ecuador. Neotropical Biodiversity 2021, 7 (1), 170–183. 10.1080/23766808.2021.1920296.

(31) Bradley, I. M.; Sevillano-Rivera, M. C.; Pinto, A. J.; Guest, J. S. Impact of Solids Residence Time on Community Structure and Nutrient Dynamics of Mixed Phototrophic Wastewater Treatment Systems. Water Res. 2019, 150, 271–282. 10.1016/j.watres.2018.11.065.

(32) Jo, S.-W.; Do, J.-M.; Na, H.; Hong, J. W.; Kim, I.-S.; Yoon, H.-S. Assessment of Biomass Potentials of Microalgal Communities in Open Pond Raceways Using Mass Cultivation. PeerJ 2020, 8, e9418. 10.7717/peerj.9418.

(33) Ma, Y.; Li, P.; Zhang, Y.; Guo, X.; Song, Y.; Zhang, Y.; Guo, Q.; Li, H.; Wang, Y.; Wan, J. Characteristics and Performance of Algal–Bacterial Granular Sludge in Photo-Sequencing Batch Reactors under Various Substrate Loading Rates. J. Environ. Manage. 2024, 368, 122216. 10.1016/j.jenvman.2024.122216.

(34) Aparicio, S.; Ríos-Mejía, A.; Gallardo-Mejías, J. P.; Robles, Á.; Borrás, L. Microalgae-Bacteria Consortia Dynamics in a Long Term Operated Membrane-Coupled High-Rate Algal Pond (MHRAP). J. Environ. Manage. 2024, 371, 123186. 10.1016/j.jenvman.2024.123186.

(35) Tedersoo, L.; Albertsen, M.; Anslan, S.; Callahan, B. Perspectives and Benefits of High-Throughput Long-Read Sequencing in Microbial Ecology. Appl. Environ. Microbiol. 2021, 87 (17), e00626–21. 10.1128/AEM.00626-21.

(36) Latz, M. A. C.; Grujcic, V.; Brugel, S.; Lycken, J.; John, U.; Karlson, B.; Andersson, A.; Andersson, A. F. Short- and Long-Read Metabarcoding of the Eukaryotic rRNA Operon: Evaluation of Primers and Comparison to Shotgun Metagenomics Sequencing. Mol. Ecol. Resour. 2022, 22 (6), 2304–2318. 10.1111/1755-0998.13623.

(37) Hugerth, L. W.; Muller, E. E. L.; Hu, Y. O. O.; Lebrun, L. A. M.; Roume, H.; Lundin, D.; Wilmes, P.; Andersson, A. F. Systematic Design of 18S rRNA Gene Primers for Determining Eukaryotic Diversity in Microbial Consortia. PLOS One 2014, 9 (4), e95567. 10.1371/journal.pone.0095567.

(38) Agustinho, D. P.; Fu, Y.; Menon, V. K.; Metcalf, G. A.; Treangen, T. J.; Sedlazeck, F. J. Unveiling Microbial Diversity: Harnessing Long-Read Sequencing Technology. Nat. Methods 2024, 21 (6), 954–966. 10.1038/s41592-024-02262-1.

(39) Greay, T. L.; Gofton, A. W.; Zahedi, A.; Paparini, A.; Linge, K. L.; Joll, C. A.; Ryan, U. M. Evaluation of 16S Next-Generation Sequencing of Hypervariable Region 4 in Wastewater Samples: An Unsuitable Approach for Bacterial Enteric Pathogen Identification. Sci. Total Environ. 2019, 670, 1111–1124. 10.1016/j.scitotenv.2019.03.278.

(40) Gehrig, J. L.; Portik, D. M.; Driscoll, M. D.; Jackson, E.; Chakraborty, S.; Gratalo, D.; Ashby, M.; Valladares, R. Finding the Right Fit: Evaluation of Short-Read and Long-Read Sequencing Approaches to Maximize the Utility of Clinical Microbiome Data. *Microb*. Genomics 2022, 8 (3), 794–812. 10.1099/mgen.0.000794.

(41) Rowan, R.; Powers, D. Molecular Genetic Identification of Symbiotic Dinoflagellates (Zooxanthellae). Mar. Ecol. Prog. Ser. 1991, 71, 65–73. 10.3354/meps071065.

(42) Medlin, L.; Elwood, H. J.; Stickel, S.; Sogin, M. L. The Characterization of Enzymatically Amplified Eukaryotic 16S-like rRNA-Coding Regions. Gene 1988, 71 (2), 491–499. 10.1016/0378-1119(88)90066-2.

(43) Quast, C.; Pruesse, E.; Yilmaz, P.; Gerken, J.; Schweer, T.; Yarza, P.; Peplies, J.; Glöckner, F. O. The SILVA Ribosomal RNA Gene Database Project: Improved Data Processing and Web-Based Tools. Nucleic Acids Res. 2012, 41 (D1), D590–D596. 10.1093/nar/gks1219.

(44) Guillou, L.; Bachar, D.; Audic, S.; Bass, D.; Berney, C.; Bittner, L.; Boutte, C.; Burgaud, G.; De Vargas, C.; Decelle, J.; Del Campo, J.; Dolan, J. R.; Dunthorn, M.; Edvardsen, B.; Holzmann, M.; Kooistra, W. H. C. F.; Lara, E.; Le Bescot, N.; Logares, R.; Mahé, F.; Massana, R.; Montresor, M.; Morard, R.; Not, F.; Pawlowski, J.; Probert, I.; Sauvadet, A.-L.; Siano, R.; Stoeck, T.; Vaulot, D.; Zimmermann, P.; Christen, R. The Protist Ribosomal Reference Database (PR2): A Catalog of Unicellular Eukaryote Small Sub-Unit rRNA Sequences with Curated Taxonomy. Nucleic Acids Res. 2012, 41 (D1), D597–D604. 10.1093/nar/gks1160.

(45) Slater, G. S. C.; Birney, E. Automated Generation of Heuristics for Biological Sequence Comparison. BMC Bioinf. 2005, 6 (1), 31–41. 10.1186/1471-2105-6-31.

(46) Robeson, M. S.; O’Rourke, D. R.; Kaehler, B. D.; Ziemski, M.; Dillon, M. R.; Foster, J. T.; Bokulich, N. A. RESCRIPt: Reproducible Sequence Taxonomy Reference Database Management. PLOS Comput. Biol. 2021, 17 (11), e1009581. 10.1371/journal.pcbi.1009581.

(47) Bolyen, E.; Rideout, J. R.; Dillon, M. R.; Bokulich, N. A.; Abnet, C. C.; Al-Ghalith, G. A.; Alexander, H.; Alm, E. J.; Arumugam, M.; Asnicar, F.; Bai, Y.; Bisanz, J. E.; Bittinger, K.; Brejnrod, A.; Brislawn, C. J.; Brown, C. T.; Callahan, B. J.; Caraballo-Rodríguez, A. M.; Chase, J.; Cope, E. K.; Da Silva, R.; Diener, C.; Dorrestein, P. C.; Douglas, G. M.; Durall, D. M.; Duvallet, C.; Edwardson, C. F.; Ernst, M.; Estaki, M.; Fouquier, J.; Gauglitz, J. M.; Gibbons, S. M.; Gibson, D. L.; Gonzalez, A.; Gorlick, K.; Guo, J.; Hillmann, B.; Holmes, S.; Holste, H.; Huttenhower, C.; Huttley, G. A.; Janssen, S.; Jarmusch, A. K.; Jiang, L.; Kaehler, B. D.; Kang, K. B.; Keefe, C. R.; Keim, P.; Kelley, S. T.; Knights, D.; Koester, I.; Kosciolek, T.; Kreps, J.; Langille, M. G. I.; Lee, J.; Ley, R.; Liu, Y.-X.; Loftfield, E.; Lozupone, C.; Maher, M.; Marotz, C.; Martin, B. D.; McDonald, D.; McIver, L. J.; Melnik, A. V.; Metcalf, J. L.; Morgan, S. C.; Morton, J. T.; Naimey, A. T.; Navas-Molina, J. A.; Nothias, L. F.; Orchanian, S. B.; Pearson, T.; Peoples, S. L.; Petras, D.; Preuss, M. L.; Pruesse, E.; Rasmussen, L. B.; Rivers, A.; Robeson, M. S.; Rosenthal, P.; Segata, N.; Shaffer, M.; Shiffer, A.; Sinha, R.; Song, S. J.; Spear, J. R.; Swafford, A. D.; Thompson, L. R.; Torres, P. J.; Trinh, P.; Tripathi, A.; Turnbaugh, P. J.; Ul-Hasan, S.; Van Der Hooft, J. J. J.; Vargas, F.; Vázquez-Baeza, Y.; Vogtmann, E.; Von Hippel, M.; Walters, W.; Wan, Y.; Wang, M.; Warren, J.; Weber, K. C.; Williamson, C. H. D.; Willis, A. D.; Xu, Z. Z.; Zaneveld, J. R.; Zhang, Y.; Zhu, Q.; Knight, R.; Caporaso, J. G. Reproducible, Interactive, Scalable and Extensible Microbiome Data Science Using QIIME 2. Nat. Biotechnol. 2019, 37 (8), 852–857. 10.1038/s41587-019-0209-9.

(48) Shen, W.; Le, S.; Li, Y.; Hu, F. SeqKit: A Cross-Platform and Ultrafast Toolkit for FASTA/Q File Manipulation. PLOS One 2016, 11 (10), e0163962. 10.1371/journal.pone.0163962.

(49) Shen, W.; Sipos, B.; Zhao, L. SeqKit2: A Swiss Army Knife for Sequence and Alignment Processing. iMeta 2024, 3 (3), e191. 10.1002/imt2.191.

(50) Weinstein, M. M.; Prem, A.; Jin, M.; Tang, S.; Bhasin, J. M. FIGARO: An Efficient and Objective Tool for Optimizing Microbiome rRNA Gene Trimming Parameters. bioRxiv April 16, 2019, p 610394. 10.1101/610394.

(51) Callahan, B. J.; McMurdie, P. J.; Rosen, M. J.; Han, A. W.; Johnson, A. J. A.; Holmes, S. P. DADA2: High-Resolution Sample Inference from Illumina Amplicon Data. Nat. Methods 2016, 13 (7), 581–583. 10.1038/nmeth.3869.

(52) Martin, M. Cutadapt Removes Adapter Sequences from High-Throughput Sequencing Reads. EMBnet.j. 2011, 17 (1), 10. 10.14806/ej.17.1.200.

(53) R Core Team. R: A Language and Environment for Statistical Computing, 2021. https://www.R-project.org/.

(54) RStudio Team. RStudio: Integrated Development Environment for R; RStudio, PBC: Boston, MA, 2022.

(55) Barrett, T.; Dowle, M.; Srinivasan, A.; Gorecki, J.; Chirico, M.; Hocking, T.; Schwendinger, B.; Krylov, I. *Data.*Table: Extension of ‘data.Frame’; manual; 2026. 10.32614/CRAN.package.data.table.

(56) Wickham, H.; Averick, M.; Bryan, J.; Chang, W.; McGowan, L. D.; François, R.; Grolemund, G.; Hayes, A.; Henry, L.; Hester, J.; Kuhn, M.; Pedersen, T. L.; Miller, E.; Bache, S. M.; Müller, K.; Ooms, J.; Robinson, D.; Seidel, D. P.; Spinu, V.; Takahashi, K.; Vaughan, D.; Wilke, C.; Woo, K.; Yutani, H. Welcome to the tidyverse. J. Open Source Software 2019, 4 (43), 1686. 10.21105/joss.01686.

(57) McMurdie, P. J.; Holmes, S. Phyloseq: An R Package for Reproducible Interactive Analysis and Graphics of Microbiome Census Data. PLOS One 2013, 8 (4), e61217. 10.1371/journal.pone.0061217.

(58) Oksanen, J.; Simpson, G. L.; Blanchet, F. G.; Kindt, R.; Legendre, P.; Minchin, P. R.; O’Hara, R. B.; Solymos, P.; Stevens, M. H. H.; Szoecs, E.; Wagner, H.; Barbour, M.; Bedward, M.; Bolker, B.; Borcard, D.; Borman, T.; Carvalho, G.; Chirico, M.; De Caceres, M.; Durand, S.; Evangelista, H. B. A.; FitzJohn, R.; Friendly, M.; Furneaux, B.; Hannigan, G.; Hill, M. O.; Lahti, L.; Martino, C.; McGlinn, D.; Ouellette, M.-H.; Ribeiro Cunha, E.; Smith, T.; Stier, A.; Ter Braak, C. J. F.; Weedon, J. Vegan: Community Ecology Package; manual; 2026. 10.32614/CRAN.package.vegan.

(59) Mariadassou, M. Phyloseq.Extended: PhyloseqExtended; manual; 2026. https://github.com/mahendra-mariadassou/phyloseq-extended.

(60) Ullrich, K. K. MSA2dist: MSA2dist Calculates Pairwise Distances between All Sequences of a DNAStringSet or a AAStringSet Using a Custom Score Matrix and Conducts Codon Based Analysis; manual; 2025. 10.18129/B9.bioc.MSA2dist.

(61) Paradis, E.; Schliep, K. Ape 5.0: An Environment for Modern Phylogenetics and Evolutionary Analyses in R. Bioinformatics 2019, 35 (3), 526–528. 10.1093/bioinformatics/bty633.

(62) Fernandes, A. D.; Macklaim, J. M.; Linn, T. G.; Reid, G.; Gloor, G. B. ANOVA-like Differential Expression (ALDEx) Analysis for Mixed Population RNA-Seq. PLOS One 2013, 8 (7), e67019. 10.1371/journal.pone.0067019.

(63) Fernandes, A. D.; Reid, J. N.; Macklaim, J. M.; McMurrough, T. A.; Edgell, D. R.; Gloor, G. B. Unifying the Analysis of High-Throughput Sequencing Datasets: Characterizing RNA-Seq, 16S rRNA Gene Sequencing and Selective Growth Experiments by Compositional Data Analysis. Microbiome 2014, 2 (1), 15–27. 10.1186/2049-2618-2-15.

(64) Gloor, G. B.; Macklaim, J. M.; Fernandes, A. D. Displaying Variation in Large Datasets: Plotting a Visual Summary of Effect Sizes. J. Comput. Graphical Stat. 2016, 25 (3), 971–979. 10.1080/10618600.2015.1131161.

(65) Wickham, H. Ggplot2: Elegant Graphics for Data Analysis; Springer-Verlag New York, 2016.

(66) Kassambara, A. Ggpubr: “ggplot2” Based Publication Ready Plots; manual; 2020. https://CRAN.R-project.org/package=ggpubr.

(67) Tiedemann, F. Gghalves: Compose Half-Half Plots Using Your Favourite Geoms; manual; 2026. https://github.com/erocoar/gghalves.

(68) Klindworth, A.; Pruesse, E.; Schweer, T.; Peplies, J.; Quast, C.; Horn, M.; Glöckner, F. O. Evaluation of General 16S Ribosomal RNA Gene PCR Primers for Classical and Next-Generation Sequencing-Based Diversity Studies. Nucleic Acids Res. 2013, 41 (1), e1–e1. 10.1093/nar/gks808.

(69) Hadziavdic, K.; Lekang, K.; Lanzen, A.; Jonassen, I.; Thompson, E. M.; Troedsson, C. Characterization of the 18S rRNA Gene for Designing Universal Eukaryote Specific Primers. PLOS One 2014, 9 (2), e87624. 10.1371/journal.pone.0087624.

(70) Parks, D. H.; Chuvochina, M.; Waite, D. W.; Rinke, C.; Skarshewski, A.; Chaumeil, P.-A.; Hugenholtz, P. A Standardized Bacterial Taxonomy Based on Genome Phylogeny Substantially Revises the Tree of Life. Nat. Biotechnol. 2018, 36 (10), 996–1004. 10.1038/nbt.4229.

(71) Bridge, P. D.; Roberts, P. J.; Spooner, B. M.; Panchal, G. On the Unreliability of Published DNA Sequences. New Phytol. 2003, 160 (1), 43–48. 10.1046/j.1469-8137.2003.00861.x.

(72) Nilsson, R. H.; Ryberg, M.; Kristiansson, E.; Abarenkov, K.; Larsson, K.-H.; Kõljalg, U. Taxonomic Reliability of DNA Sequences in Public Sequence Databases: A Fungal Perspective. PLOS One 2006, 1 (1), e59. 10.1371/journal.pone.0000059.

(73) Kozlov, A. M.; Zhang, J.; Yilmaz, P.; Glöckner, F. O.; Stamatakis, A. Phylogeny-Aware Identification and Correction of Taxonomically Mislabeled Sequences. Nucleic Acids Res. 2016, 44 (11), 5022–5033. 10.1093/nar/gkw396.

(74) Edgar, R. Taxonomy Annotation and Guide Tree Errors in 16S rRNA Databases. PeerJ 2018, 6, e5030. 10.7717/peerj.5030.

(75) Mondal, S.; Bera, S.; Mishra, R.; Roy, S. Redefining the Role of Microalgae in Industrial Wastewater Remediation. Energy Nexus 2022, 6, 100088. 10.1016/j.nexus.2022.100088.

(76) Chokshi, K.; Pancha, I.; Ghosh, A.; Mishra, S. Microalgal Biomass Generation by Phycoremediation of Dairy Industry Wastewater: An Integrated Approach towards Sustainable Biofuel Production. Bioresour. Technol. 2016, 221, 455–460. 10.1016/j.biortech.2016.09.070.

(77) Cheng, H.; Liu, Q.; Zhao, G.; Yu, S.; Li, L. The Comparison of Three Common Microalgae for Treating Piggery Wastewater. Desalin. Water Treat. 2017, 98, 59–65. 10.5004/dwt.2017.21613.

(78) Pandey, A.; Srivastava, S.; Kumar, S. Sequential Optimization of Essential Nutrients Addition in Simulated Dairy Effluent for Improved Scenedesmus Sp ASK22 Growth, Lipid Production and Nutrients Removal. Biomass Bioenergy 2019, 128, 105319. 10.1016/j.biombioe.2019.105319.

(79) Ansari, F. A.; Singh, P.; Guldhe, A.; Bux, F. Microalgal Cultivation Using Aquaculture Wastewater: Integrated Biomass Generation and Nutrient Remediation. Algal Res. 2017, 21, 169–177. 10.1016/j.algal.2016.11.015.

(80) Han, W.; Jin, W.; Li, Z.; Wei, Y.; He, Z.; Chen, C.; Qin, C.; Chen, Y.; Tu, R.; Zhou, X. Cultivation of Microalgae for Lipid Production Using Municipal Wastewater. Process Saf. Environ. Prot. 2021, 155, 155–165. 10.1016/j.psep.2021.09.014.

(81) Da Fontoura, J. T.; Rolim, G. S.; Farenzena, M.; Gutterres, M. Influence of Light Intensity and Tannery Wastewater Concentration on Biomass Production and Nutrient Removal by Microalgae Scenedesmus Sp. Process Saf. Environ. Prot. 2017, 111, 355–362. 10.1016/j.psep.2017.07.024.

(82) Wang, S.; Ortiz Tena, F.; Dey, R.; Thomsen, C.; Steinweg, C.; Kraemer, D.; Grossman, A. D.; Belete, Y. Z.; Bernstein, R.; Gross, A.; Leu, S.; Boussiba, S.; Thomsen, L.; Posten, C. Submerged Hollow-Fiber-Ultrafiltration for Harvesting Microalgae Used for Bioremediation of a Secondary Wastewater. Sep. Purif. Technol. 2022, 289, 120744. 10.1016/j.seppur.2022.120744.

(83) Gardner-Dale, D. A.; Bradley, I. M.; Guest, J. S. Influence of Solids Residence Time and Carbon Storage on Nitrogen and Phosphorus Recovery by Microalgae across Diel Cycles. Water Res. 2017, 121, 231–239. 10.1016/j.watres.2017.05.033.

(84) Gong, Y.; Patterson, D. J.; Li, Y.; Hu, Z.; Sommerfeld, M.; Chen, Y.; Hu, Q. Vernalophrys Algivore Gen. Nov., Sp. Nov. (Rhizaria: Cercozoa: Vampyrellida), a New Algal Predator Isolated from Outdoor Mass Culture of Scenedesmus Dimorphus. Appl. Environ. Microbiol. 2015, 81 (12), 3900–3913. 10.1128/AEM.00160-15.

(85) Zhang, H.; Patterson, D. J.; He, Y.; Wang, H.; Yuan, D.; Hu, Q.; Gong, Y. Kinopus Chlorellivorus Gen. Nov., Sp. Nov. (Vampyrellida, Rhizaria), a New Algivorous Protist Predator Isolated from Large-Scale Outdoor Cultures of Chlorella Sorokiniana. Appl. Environ. Microbiol. 2022, 88 (22), e01215–22. 10.1128/aem.01215-22.

(86) Woznica, A. What Choanoflagellates Can Teach Us about Symbiosis. PLOS Biol. 2024, 22 (4), e3002561. 10.1371/journal.pbio.3002561.

(87) Pohl, N.; Solbach, M. D.; Dumack, K. The Wastewater Protist Rhogostoma Minus (Thecofilosea, Rhizaria) Is Abundant, Widespread, and Hosts Legionellales. Water Res. 2021, 203, 117566. 10.1016/j.watres.2021.117566.

(88) Dumack, K.; Flues, S.; Hermanns, K.; Bonkowski, M. Rhogostomidae (Cercozoa) from Soils, Roots and Plant Leaves (Arabidopsis Thaliana): Description of Rhogostoma Epiphylla Sp. Nov. and R. Cylindrica Sp. Nov. Eur. J. Protistol. 2017, 60, 76–86. 10.1016/j.ejop.2017.06.001.

