## Supplemental Figures for "Systematic comparisons between long-read and short-read based amplicon sequencing to characterize mixed microalgal communities"

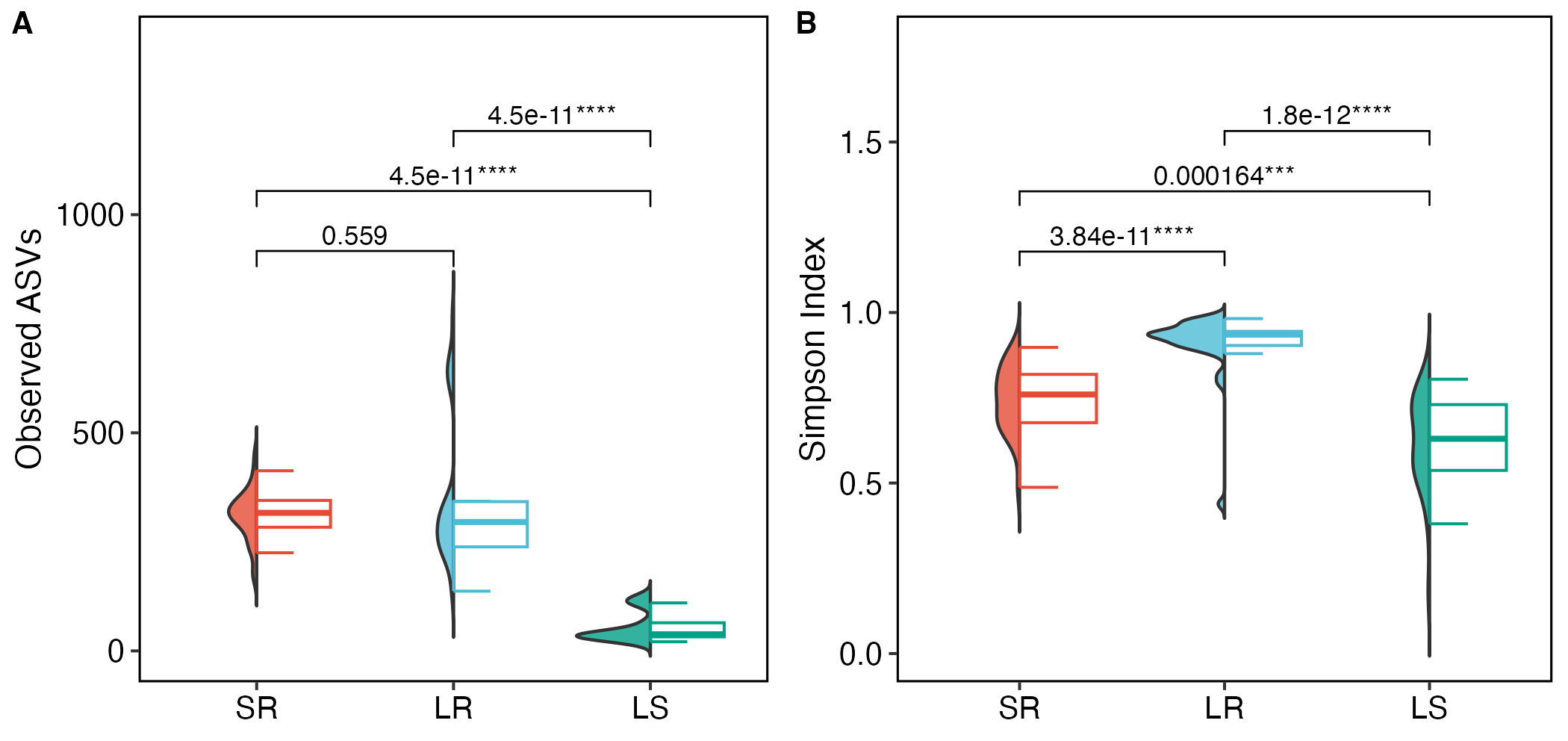

**Figure S1**: Comparison of additional alpha diversity metrics across three 18S rRNA gene sequencing datasets. (A) Observed ASVs. (B) Simpson diversity index. Half-violin plots illustrate the density distribution of each metric; half-boxplots show median and interquartile range. Pairwise differences were assessed using Wilcoxon tests with Benjamini-Hochberg correction. SR, LR, and LS datasets are shown in vermillion-red, sky cyan, and teal-green, respectively. SR: short-read V8V9 region sequenced using Illumina platform; LR: long-read full-length 18S rRNA gene sequenced using PacBio platform; LS: computationally extracted V8V9 regions from the LR dataset.

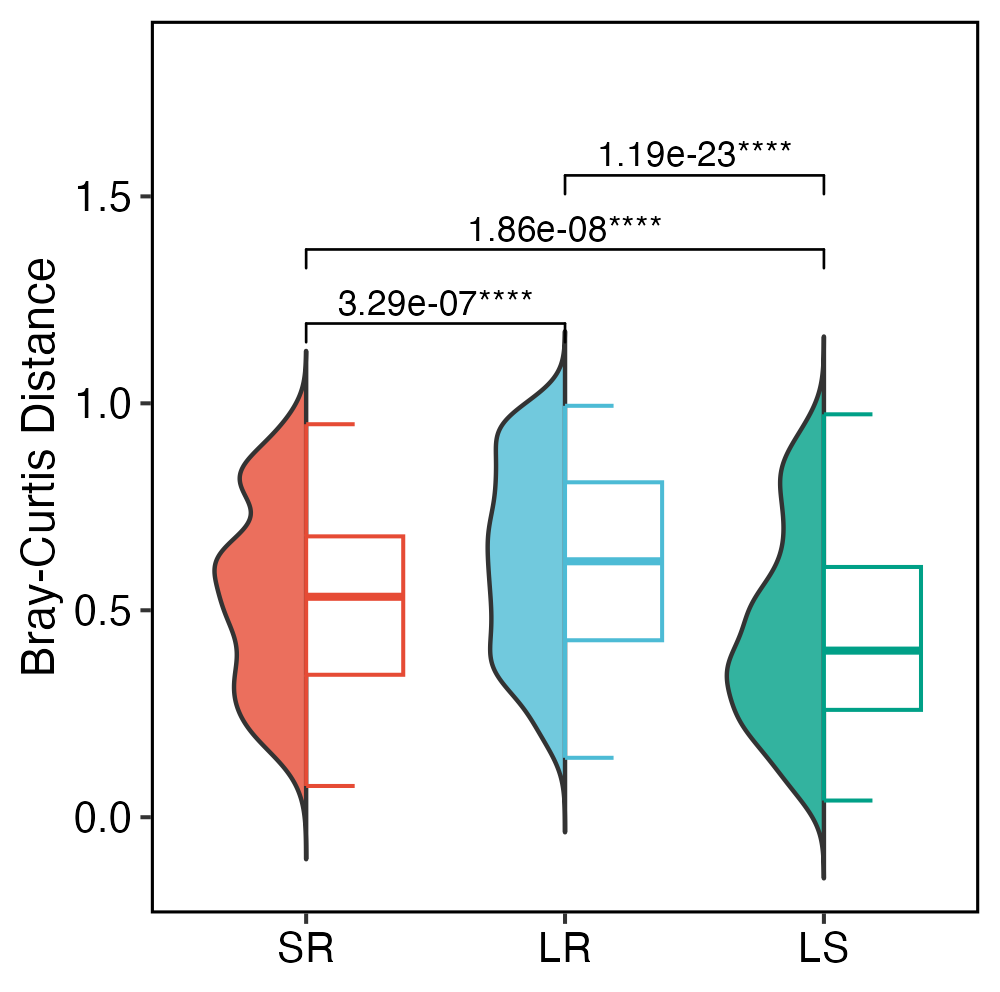

**Figure S2**: Comparison of Bray-Curtis distances across three 18S rRNA gene sequencing datasets. Half-violin plots illustrate the density distribution; half-boxplots show median and interquartile range. Pairwise differences were assessed using Wilcoxon tests with Benjamini-Hochberg correction. SR, LR, and LS datasets are shown in vermillion-red, sky cyan, and teal-green, respectively. SR: short-read V8V9 region sequenced using Illumina platform; LR: long-read full-length 18S rRNA gene sequenced using PacBio platform; LS: computationally extracted V8V9 regions from the LR dataset.

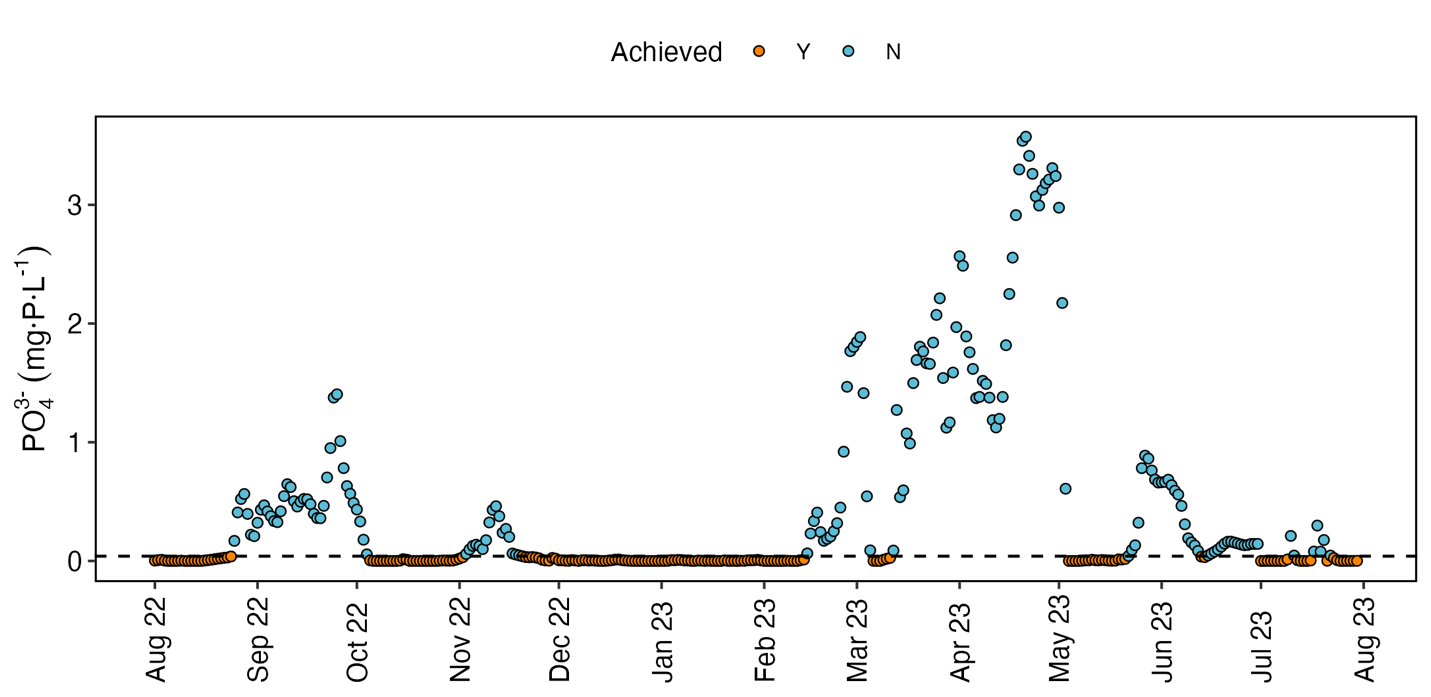

**Figure S3**: Temporal variation of daily mean effluent orthophosphate concentrations from August 1, 2022 to July 30, 2023, measured via online sensors integrated with a supervisory control and data acquisition system (SCADA) system. The dashed line indicates the phosphorus discharge limit (0.04 mg-P·L^-1^). Samples meeting (Y) and exceeding (N) the limit are shown in orange and blue, respectively. Samples are color-coded to indicate compliance with the phosphorus discharge limit (0.04 mg-P·L^−1^); orange points represent samples meeting the limit, while blue points indicate samples exceeding the threshold.
